# Transcriptomic plasticity and symbiont shuffling underpin *Pocillopora* acclimatization across heat-stress regimes in the Pacific Ocean

**DOI:** 10.1101/2021.11.12.468330

**Authors:** Eric J Armstrong, Julie Lê-Hoang, Quentin Carradec, Jean-Marc Aury, Benjamin Noel, Julie Poulain, Caroline Belser, Corinne Da Silva, Patrick Wincker, Tara Pacific Consortium

## Abstract

The characterization of adaptation and acclimation capacities of coral holobionts is crucial for anticipating the impact of global climate change on coral reefs. Understanding the extent to which the coral host and its photosymbionts contribute to adaptive and/or plastic responses in the coral metaorganism is important. In this study, we highlight new and complex links between coral genomes, transcriptomes, and environmental features in Pocilloporid corals at basin-wide scale. We analyzed metagenomic and metatranscriptomic sequence data from *Pocillopora* colonies sampled from 11 islands across the Pacific Ocean in order to investigate patterns of gene expression in both the host and photosymbiont across an environmental gradient. Single nucleotide polymorphisms (SNPs) analysis partitioned coral hosts and algal photosymbionts into five genetic lineages each. We observed strong host-symbiont fidelity across environments except at islands where recent and/or historical heat stress may have induced a symbiont shift towards more heat-tolerant lineages in some colonies. Host gene expression profiles were strongly segregated by genetic lineage and environment, and were significantly correlated with several historical sea surface temperature (SST) traits. Symbiont expression profiles were less dependent on environmental context than the host and were primarily driven by algal genotype. Overall, our results suggest a three-tiered strategy underpinning thermal acclimatization in *Pocillopora* holobionts with 1) host-photosymbiont fidelity, 2) host transcriptomic plasticity, and 3) photosymbiont shuffling playing progressive roles in response to elevated SSTs. Our data provide a reference for the biological state of coral holobionts across the Indo-Pacific and demonstrate the power of disentangling environmental and genetic effects to provide new insights into corals’ capacities for acclimatization and adaptation under environmental change.

## 1. INTRODUCTION

Coral reefs are ecologically and economically important ecosystems whose existence depends upon the mutualistic, photosymbiotic, association between certain Cnidarian hosts and their dinoflagellate symbionts. Breakdown of this association can lead to expulsion of algal symbionts and ultimately result in coral mortality and reef loss (Hoegh-Guldberg 1999; Hughes *et al.* 2017, 2018). Recent environmental perturbations, particularly rising sea surface temperatures and increasing frequency of extreme heating events (i.e., marine heatwaves), are disrupting coral-dinoflagellate photosymbioses worldwide, increasing the frequency and severity of coral bleaching events (Hughes *et al.* 2018). There is therefore rising concern about the potential for heating-driven local extirpation of coral species and severe reef loss in the coming century. However, in some regions, corals exhibit higher than average heat tolerances (Osman *et al.* 2018) or show the ability to rapidly respond to and recover from acute thermal challenges (Savary *et al.* 2021). Repeated sublethal exposure to elevated sea surface temperatures can also drive positive acclimatization (phenotypic plasticity) and/or selection for thermotolerant genotypes (i.e., local adaptation; Middlebrook *et al.* 2008; Savary *et al.* 2021). However in other species, warm preconditioning had no or negative effects on holobiont performance during subsequent thermal challenges (Middlebrook *et al.* 2012; Schoepf *et al*. 2019). The capacity for thermal acclimatization and/or adaptation may therefore play an important role in determining potential ‘winners and losers’ among coral species in the face of global change (Wright *et al.* 2019; Savary *et al.* 2021). Determining whether acclimatization capacity can keep pace with projected environmental change and in which symbiotic assemblages, is therefore of paramount importance for predicting responses of the coral-dinoflagellate photosymbiosis under a warming climate.

Adaptation to the local thermal environment has been well-documented across numerous phyla (Hereford 2009; Sanford and Kelly 2011) including hermatypic corals (Kenkel *et al.* 2013; Kenkel and Matz 2017). In some cases, symbiont shuffling from thermally-sensitive to thermally-tolerant Symbiodiniaceae confers a higher resistance of the coral holobiont to heat stress (Cunning *et al.* 2015). In other cases, rapid thermal acclimation is achieved through altered gene expression in the host and/or symbiont (i.e., transcriptomic plasticity) resulting in increased whole organism (holobiont) thermal tolerance (Kenkel and Matz 2017; Savary *et al.* 2021). Coral holobiont thermal sensitivity is therefore dependent on an array of interacting drivers, including environmental history (Middlebrook *et al.* 2008; Safaie *et al.* 2018; Schoepf *et al.* 2020; Wall *et al.* 2021; Savary *et al.* 2021), endosymbiont community composition (Hoadley *et al.* 2019; Qin *et al.* 2019; Cunning and Baker 2020; Dilworth *et al.* 2021), and host genotype (Barshis *et al.* 2013; Dilworth *et al.* 2021; Drury *et al.* 2021). However, the relative role of each of these factors in determining holobiont expression *in situ* remains poorly resolved. A remaining challenge, therefore, is to better understand how environmental context affects the complex interplay between the host and symbiont and to what extent certain genotypes and or host/symbiont pairings might confer resistance to projected environmental change.

Understanding how corals will respond to future climate change requires prior knowledge regarding their capacities for acclimatization (e.g., holobiont flexibility and transcriptomic plasticity) as well as their capacities for adaptation. The mechanisms underlying these capacities as well as their limits and extent among different coral holobiotypes and the relative roles of each symbiotic partner remain under intense investigation. In addition, holobiont transcriptomic adaptation/acclimation can be difficult to assess in some cases because gene expression can be highly variable between polyps within a single coral colony (Drake *et al.* 2021). Finally, acclimation or adaptation of a coral holobiont at the local (reef) scale is often not representative of capacities at larger (ocean) scales and therefore does not allow for accurate global projections of the survival or decline of a species or of reef ecosystems.

Given the variety of components involved in maintaining the coral-dinoflagellate mutualism against environmental perturbation, recent analyses have begun to adopt a more integrative and multidisciplinary approach to understanding the impacts of climate change on coral health. Integration of various analysis methods (genomic, transcriptomic, barcoding, imaging, etc.) is needed to draw a global picture of coral holobiont adaptation and acclimation capacities. The *Tara* Pacific expedition provides a unique opportunity to address these important questions because it allows for comparative analysis of holobiont regulation at ocean scale - covering many different local environments as well as different species (Planes et al. 2019). To this end, we investigated gene expression profiles of 102 colonies of *Pocillopora spp.* and their associated endosymbionts alongside environmental context data from 11 islands across the Pacific Ocean in order to assess the relative contributions of environmental and genetic factors in determining coral holobiont gene expression *in situ*. We used genome-wide gene expression profiling, and variation partitioning to examine the role of transcriptomic plasticity in local adaptation among various *Pocillopora* hosts and their associated dinoflagellate endosymbionts (*Symbiodiniaceae*) across environmentally distinct reef environments.

## 2. MATERIALS & METHODS

### 2.1. Site selection, coral colony sampling, and environmental metadata collection

A total of 102 *Pocillopora spp.* colonies from 11 islands (Islas de las Perlas, Coïba, Malpelo, Rapa Nui, Ducie, Gambier, Moorea, Aitutaki, Niue, Upolu, and Guam) were sampled as part of the *Tara* Pacific expedition (Planes et al. 2019). At each island, fragments of three coral colonies were collected from each of three reef sites yielding a total of ca. 9 colonies sampled per island (Table 1). Reef site sampling protocols are presented in detail in Gorsky *et al.* 2019. Fragments of sampled coral colonies were removed from the reef and brought back on board the *Tara* vessel for processing of DNA/RNA. Several environmental parameters were measured *in situ* at the time of collection and historical sea surface temperature (SST) data (including degree heating week data) for each sampling site were obtained as described by Gorsky *et al*. (2019; Table 1).

**Table 1.** Reef site collection information and top explanatory environmental context data.

| Island | Reef Site | Colonies | SST (°C) | PAR | pH | PO<sub>4</sub><sup>3-</sup> | TSA_DCW_s<br>tdlength<br>(days) | TSA_DHW_I<br>astrecovery<br>(days) | SST_anomal<br>y_frequency<br>_std (days) |
| --- | --- | --- | --- | --- | --- | --- | --- | --- | --- |
| Islas de las<br>Perlas | Site 1 | 3 | --- | --- | --- | --- | 21.21 | 245 | 22.79 |
|  | Site 2 | 3 | 28.95 | 41.19 | 8.03 | 0.10 | 23.33 | 280 | 23.56 |
|  | Site 3 | 2 | 28.55 | 36.43 | 8.03 | 0.09 | 13.44 | 212 | 20.58 |
| Coiba | Site 1 | 3 | --- | --- | --- | --- | 48.27 | 92 | 40.79 |
|  | Site 2 | 3 | --- | --- | --- | --- | 43.16 | 78 | 41.28 |
|  | Site 3 | 3 | 28.28 | 49.71 | 8.04 | 0.15 | 41.35 | 65 | 36.95 |
| Malpelo | Site 1 | 10 | --- | --- | --- | --- | 23.87 | 348 | 33.77 |
| Rapa Nui | Site 1 | 3 | 20.81 | 38.23 | 8.07 | 0.07 | 20.41 | 499 | 17.80 |
|  | Site 2 | 3 | 20.74 | 38.23 | 8.07 | 0.07 | 19.59 | 3816 | 18.83 |
|  | Site 3 | 3 | --- | --- | --- | --- | 24.12 | 3805 | 14.93 |
|  | Site 4 | 3 | 20.74 | 36.25 | 8.07 | 0.08 | 19.32 | 5239 | 18.15 |
| Ducie | Site 1 | 3 | 21.90 | 43.11 | 8.07 | 0.06 | 17.53 | 3778 | 22.37 |
|  | Site 2 | 3 | 21.90 | 42.89 | 8.07 | 0.06 | 20.44 | 3780 | 22.01 |
|  | Site 3 | 3 | 21.81 | 42.89 | 8.07 | 0.06 | 22.03 | 3780 | 22.06 |
| Gambier | Site 1 | 3 | 23.03 | 49.88 | 8.07 | 0.07 | 12.92 | 126 | 16.76 |
|  | Site 2 | 3 | 23.02 | 50.30 | 8.07 | 0.06 | 22.19 | 133 | 16.92 |
|  | Site 3 | 3 | 23.15 | 49.59 | 8.07 | 0.06 | 28.86 | 130 | 19.56 |
| Moorea | Site 1 | 3 | 28.67 | 37.11 | 8.06 | 0.17 | 27.76 | 172 | 7.31 |
|  | Site 2 | 3 | 28.51 | 38.14 | 8.06 | 0.18 | 36.63 | 173 | 7.40 |
|  | Site 3 | 3 | 28.64 | 41.25 | 8.04 | 0.17 | 26.39 | 170 | 8.35 |
| Aitutaki | Site 1 | 3 | 27.35 | 46.18 | 8.02 | 0.17 | 33.71 | 3832 | 10.01 |
|  | Site 2 | 3 | 27.48 | 47.93 | 8.03 | 0.17 | 30.24 | 3836 | 11.25 |
|  | Site 3 | 3 | 27.28 | 46.18 | 8.02 | 0.18 | 25.00 | 3834 | 10.70 |
| Niue | Site 1 | 3 | 28.32 | 45.21 | 8.04 | 0.14 | 31.50 | 5315 | 14.84 |
|  | Site 2 | 3 | 28.40 | 48.58 | 8.02 | 0.15 | 35.37 | 961 | 14.10 |
|  | Site 3 | 3 | 28.28 | 45.90 | 8.04 | 0.13 | 39.38 | 2759 | 23.09 |
| Upolu | Site 1 | 3 | 31.08 | 47.41 | 7.96 | 0.22 | 32.79 | 558 | 8.36 |
|  | Site 2 | 3 | 29.59 | 57.85 | 7.99 | 0.22 | 33.11 | 566 | 10.59 |
|  | Site 3 | 3 | 30.40 | 54.40 | 7.97 | 0.23 | 28.39 | 579 | 11.38 |
| Guam | Site 1 | 3 | 28.16 | 43.38 | 8.02 | 0.06 | 10.72 | 1195 | 8.67 |
|  | Site 2 | 3 | 28.13 | 41.67 | 8.02 | 0.07 | 21.78 | 5383 | 10.88 |
|  | Site 3 | 3 | 28.10 | 39.01 | 8.01 | 0.06 | 34.50 | 5384 | 10.36 |

SST, PAR, pH, PO<sub>4</sub><sup>3-</sup>: mean sea surface temperature, photosynthetically active radiation, seawater pH, and concentration of phosphate at the time of sampling. PAR was estimated from satellite data and all variables are averaged across two colonies per site.

### 2.2. DNA/RNA isolation, library preparation, and sequencing

Coral fragments were processed to extract and isolate DNA/RNA as described by Belser *et al.* (*in prep*).

### 2.3. Host/Symbiodiniaceae diversity and Cladocopium population structure

#### 2.3.1. Host lineage assignation

We performed host lineage assignation as described by Armstrong *et al.* (*in prep*; Figure S1). Briefly, we identified a set of genome-wide single nucleotide polymorphisms (SNPs) from metagenomic reads mapped to the *Pocillopora meandrina* genomic reference (Aury *et al. in prep*) using the Genome Analysis Toolkit program (GATK, v3.7.0; Van der Auwera and O’Connor 2020). We followed the best practices guide for variant discovery with GATK which included indexing of the genomic reference (picardtools v2.6.0, CreateSequenceDictionary), followed by identification of realignment targets (GATK RealignerTargetCreator) and realignment around detected indels (GATK, IndelRealigner). Variants were called for each colony individually (GATK, HaplotypeCaller) and resulting variant call files (VCFs) were merged into island-specific cohorts (GATK, CombineGVCFs) before performing joint genotyping across all 11 islands (GATK, GenotypeGVCFs) with polyploidy defined at 1 (Poplin *et al.* 2018). The resulting SNPs were filtered using VCFtools (v0.1.12; Danecek *et al.* 2011) to include only biallelic sites with minor allele frequencies ≥ 0.05, quality scores ≥ 30, and no missing data across colonies. These curated SNPs were then used to identify divergent host lineages using a coalescent analysis in RaxML.

#### 2.3.2. Cladocopium lineage assignation

To determine which *Symbiodiniaceae* lineages were present in each coral colony we first used PCR-amplified ITS2 reads sequenced for each sample. ITS2 profiles were obtained with the SymPortal method (Hume *et al.* 2019) which uses the intragenomic diversity of *Symbiodiniaceae* ITS2 to define ITS2 type profiles based on consistent co-occurrence of intragenomic ITS2 variants across all samples. After assigning ITS2 profiles to each colony, we further analyzed the 83 coral colonies containing only a single *Cladocopium* ITS2 profile in order to determine their population structure across the sampled islands. SNPs from these 83 colonies were identified from metatranscriptomic reads previously aligned on the predicted coding sequences of *Cladocopium goreaui* genome (see next paragraph) following the same best practices protocol as described for the host above. SNPs were filtered using VCFtools (v0.1.12; Danecek *et al.* 2011) to include only biallelic SNP with a quality score ≥ 30 and a coverage ≥ 4 in the 83 samples. The frequencies of the 3,712 SNP were clusterized using the Hclust function in R with Complete Linkage clustering method.

### 2.4. Gene expression levels of Pocillopora and Cladocopium from metatranscriptomic reads

Metatranscriptomic reads (Illumina-generated 150-bp PE) were aligned to predicted coding sequences of the *Pocillopora meandrina* host reference genome (Aury *et al. in prep*) as well as to predicted coding sequences of the *Cladocopium goreaui* genome (Liu et al. 2018b) and *Durusdinium* D2 transcriptome (Ladner *et al.* 2012) using Burrows–Wheeler Transform Aligner (BWA-mem, v0.7.15) with the default settings (Li and Durbin 2009). Host- and symbiont-mapped reads were then sorted and processed using SAMtools v1.10.2 (Li *et al.* 2009) to generate respective bam files. A read was considered a host contig if its sequence aligned to the *P. meandrina* predicted coding sequence with ≥ 95% of sequence identity and with ≥ 50% of the sequence aligned. *Cladocopium* reads aligned with *Cladocopium goreaui* coding sequences and *Durusdinium D2* transcriptome were mapped against each of the symbiont databases and assigned in the same manner as described above except with a more stringent assignation cutoff of ≥ 98% of sequence identity and with ≥ 80% of the read length of the sequence aligned. All assigned reads were further filtered to remove sequences which contained > 75% of low-complexity bases and < 30% high-complexity bases and resulting read alignments were visualized using the Integrated Genomics Viewer (Robinson 2011). Read counts were normalized as transcript per million (TPM).

### 2.5. Bioinformatic analyses

To evaluate the relative influence of the environment and genotype on coral host and *Cladocopium* expression profiles we used three complementary approaches, namely (1) quantitative partitioning of gene expression variance into the fraction attributable to each driver using a linear mixed model approach (variation partitioning), (2) discriminant analysis of principal components (DAPC), and (3) constrained correspondence analysis informed by both historical and *in situ* environmental drivers. We also performed a phylogenetically-informed ANOVA to identify genes whose expression variation was linked to lineage divergence within Pocillopora and Cladocopium C1 and which therefore represent potential adaptive loci between lineages.

#### 2.5.1. Quantitative partitioning of gene expression variance

In the first approach, we used the variancePartition package in R (v 1.21.2; Hoffman and Schadt 2016; Hoffman and Roussos 2021) to partition the variance attributable to the environment, the host genotype, and the symbiont genotype in the *Pocillopora* and *Cladocopium* expression datasets. Prior to performing the variation partitioning analysis, a correction was applied to remove a bias due to the two different library preparation methods used for RNA sequencing following the procedure described by Hoffman *et al.* Three variables were defined as random effects: *Pocillopora* genetic lineage, *Cladocopium* genetic lineage, and the sampling island. To take into account the sampling effect on the variance for each gene we repeated the variation partitioning analysis 100 times with a random removal of two samples.

Genes with a median explained variance > 50% for one of the three selected variables across these 100 rounds were selected for downstream analysis. For each selected gene, a z-score (the number of standard deviations from the mean expression of all samples) was calculated for each island or genetic clade. A z-score > 0.5 or < -0.5 was used to consider the gene as over or under-expressed respectively. These differentially expressed genes across all samples, hereafter referred to as top variant genes, were retained for functional enrichment analyses and were represented with the upsetR package (Conway *et al.* 2017).

#### 2.5.2. Discriminant analysis of principal components

In the first approach, we used the DAPC package in R (Jombart 2008; Jombart *et al.* 2010) to determine which genes are best able to discriminate between pre-defined groups of samples (i.e., based on either their genetic lineage or on their environment of collection) and to assess how well these groups could be distinguished from one another based on similarity among their expression profiles. The purpose of DAPC is to find the linear combinations of genes which maximize the differences between pre-specified groups while minimizing the within-group variance thereby allowing for the determination of which genes most strongly discriminate between the pre-defined groups. Coefficients of these genes are called loadings and higher loading scores indicate a stronger discriminatory ability. The linear combinations of these genes’ expression values are referred to as discriminant functions and serve to orient the groups in multi-dimensional space.

We performed three discriminant analyses each for the host and algal symbiont with the first model using the island of collection (i.e., the environmental effect) as the pre-defined grouping factor, the second using coral genetic lineage and photosymbiont genetic lineage as the grouping factor for the coral and photosymbiont, respectively, (i.e., the primary genetic effect), and the third using the genetic lineage of the symbiont (photosymbiont for the host and vice versa) as the grouping variable (i.e., the symbiotic partner effect). The input for these analyses were the normalized expression data (i.e., variance stabilized transformed counts) for all genes with 10 or more counts in at least 90% of colonies (host n = 28,732; symbiont n = 20,338). Firstly, we performed an ordination analysis to extract principal Components (PCs). We then computed a-scores to determine the optimal number of PCs to retain under each clustering model for subsequent cluster identification using discriminant analysis (Jombart et al. 2010). A-score optimization resulted in our selecting to retain 12 PCs for the host (for all three models) and 11 PCs for the algal symbiont (all models). Group memberships were then independently predicted for the colonies based on DAPC scores.

Proportions of colonies correctly reassigned to their pre-defined groups, PCs and discriminant functions (DF) retained, and the overall proportion of variance explained by each of the clustering models are included in Table S9. The relative strength of environmental-, genetic-, and symbiont-impacts on *Pocillopora* and *Cladocopium* gene expression was assessed by comparing the proportion of colonies that were correctly re-assigned to their pre-defined groups under each clustering model. A higher proportion of colonies correctly reassigned to their pre-defined groups indicates greater power to discriminate between divergent expression profiles and, therefore, a greater influence of that grouping factor on gene expression. In addition to this assessment of grouping-factor influence we also identified genes whose expression profiles most strongly contributed to model discriminant axes. Genes with discriminant axis loading scores within the upper quartile of scores (i.e., top 25%) were considered as significant contributors to that axis, are referred to hereafter as discriminant genes, and were retained for analysis of functional enrichments. Because there was strong host-symbiont fidelity across all lineages, there was consequently significant overlap between discriminant genes recovered from the primary genetic and symbiotic partner DAPC models. We therefore further divided these discriminant genes into two subcategories: genes which were shared between the two models (shared discriminant genes) and genes which were unique to one of the two models (unique discriminant genes). We use these subcategories to distinguish between analyses of all discriminant genes and unique discriminant genes, respectively, throughout this manuscript.

#### 2.5.3. Expression variance and evolution model (EVE)

The combination of RNA-seq and genomic datasets in this study allowed us to test whether variation in gene expression differed significantly among species (i.e., expression divergence) and/or within species across environments (i.e., expression diversity or plasticity). By treating gene expression levels as quantitative traits, we used the expression variance and evolution model (EVE) to parameterize the ratio (β) of population (i.e., within-species) to evolutionary (i.e., among-species) expression variance (Rohlfs and Nielsen 2015; Avila-Magaña et al. 2021). This, in turn, allowed us to detect genes which may be under selection or which exhibit high plasticity across environments.

In brief, the EVE model works by identifying genes whose β-values fall significantly above or below a given threshold established from genes which are not under strong selection. In the presence of stabilizing or no selection, the β-value for a given gene will remain constant and this constant forms the threshold for identifying β-values that deviate from the ‘no selection’ assumption. However, if the variation in gene expression within-species is significantly greater than that among-species (i.e., a high β-value) this indicates that that gene shows high *diversity* (phenotypic plasticity) in gene expression within a lineage and therefore is identified as a potential plastic candidate gene. Conversely, when β is low, indicating a higher among-species expression variation than within-species variance, this indicates that that gene shows lineage-specific *divergence* in expression and may therefore represent a potential adaptive candidate gene. The EVE model uses a likelihood ratio test (LRT) to examine whether the β-value for a given gene deviates significantly from the χ2 distribution of all tested genes which is expected under the null model. We identified genes with a β-value greater/lower than expected at a cutoff value of FDR < 0.1 as potential plastic/adaptive candidate genes in both *Pocillopora* and *Cladocopium C1*. These genes were retained for functional enrichment analyses and are hereafter referred to as diverse and divergent genes, respectively.

#### 2.5.4. Constrained correspondence analysis of environmental data

We also performed a constrained correspondence analysis (CCA) between the host/symbiont expression data and the historical SST data in order to visualize how expression data were structured with respect to potential environmental drivers. In order to avoid redundant environmental data we first identified highly correlated variables (Pearson R^2^ ≥ 0.70) and kept only one of these for further analysis. We then used the bioenv function of the Vegan package in R (v2.5.6) to identify the best subset of environmental variables so that the Euclidean distances of the scaled environmental variables would have the maximum correlation with expression profile dissimilarities (Clarke and Ainsworth 1993). Permutation testing was then implemented in order to select the top environmental variables (i.e., those with the highest explanatory power at p ≤ 0.01) driving differentiation in overall gene expression profiles across the colonies sampled.

#### 2.5.5. Analysis of functional enrichments among genes of interest

Functional annotations for predicted coding sequences of the *Pocillopora* host genome were extracted from the genomic reference as described by Aury *et al.* (*in prep*). These included Interproscan (IPR) and Pfam protein family terms as well as corresponding Gene Ontology (GO) annotations attributed with the Interpro2GO correspondence table (version 2020/06/13; Mitchell et al. 2015). Functional annotation of *Cladocopium goreaui* genes were recovered from the published genome (Liu et al. 2018b).

Functional enrichment among top variant (variation partitioning) and discriminant (DAPC) genes were analyzed in the host using the GOSeq (v1.40.0; Young *et al.* 2010) and GO_MWU (Wright *et al.* 2015) packages in R. For top variant genes, the number of GO annotations assigned to genes within that interest group was compared to the number of annotations assigned to the rest of the dataset, to evaluate whether any ontologies were more highly represented within the module than expected by chance (i.e., Fisher’s exact test in GOSEq). However, for discriminant genes, because an additional quantitative measure of gene importance (i.e., loading score) was available for each gene, we performed a Mann-Whitney U Test (GO_MWU package) to identify enriched GO terms. Loading scores for each discriminant function were normalized to the highest score for that function yielding values ranging from 0 (no significant contribution) to 1 (highest contribution). We then initiated the gomwuStats function of the GO_MWU package in module mode and the normalized loading scores used in place of the module-membership (kME) values in the Mann-Whitney U Test.

*Cladocopium* functional enrichment analyses were performed using independent Fisher’s exact tests for each Pfam (protein family) domain between genes present in each gene-of-interest category (top variant, discriminant, etc.) as compared to all expressed genes of *Cladocopium*. P-values were corrected for multiple testing with the Benjamini & Hochberg method implemented in the p.adjust function in R. For each Pfam term, the number of annotations assigned to genes within an interest group (i.e., top variant and discriminant genes) was compared to the number of annotations assigned to the genome, to evaluate whether any ontologies were more highly represented within the module than expected by chance (Fisher’s exact test).

## 3. RESULTS

### 3.1. Lineage assignation and characterization of host-symbiont specificity

Among the 102 colonies sampled across the 11 islands (Figure 1A), we identified five distinct *Pocillopora* host lineages, hereafter named SVD1 to SVD5 and observed a strong geographic partitioning of host lineages across the Pacific basin (Figure 1B). *Pocillopora* SVD 4 was the sole species present among colonies sampled from the eastern Pacific (Las Perlas, Coïba, and Malpelo) and was also present on Ducie and Gambier. Host SVD 5 was the only species sampled on Rapa Nui and was also present on Ducie and Moorea. *Pocillopora* SVD1 was restricted to the central Pacific (Ducie, Moorea, and Gambier) whereas *Pocillopora* SVD2 and SVD3 were both present in the western Pacific (Aitutaki, Niue, Upolu, and Guam). The SVD2 lineage was also present on the island of Moorea in the central Pacific.

**Figure 1.**
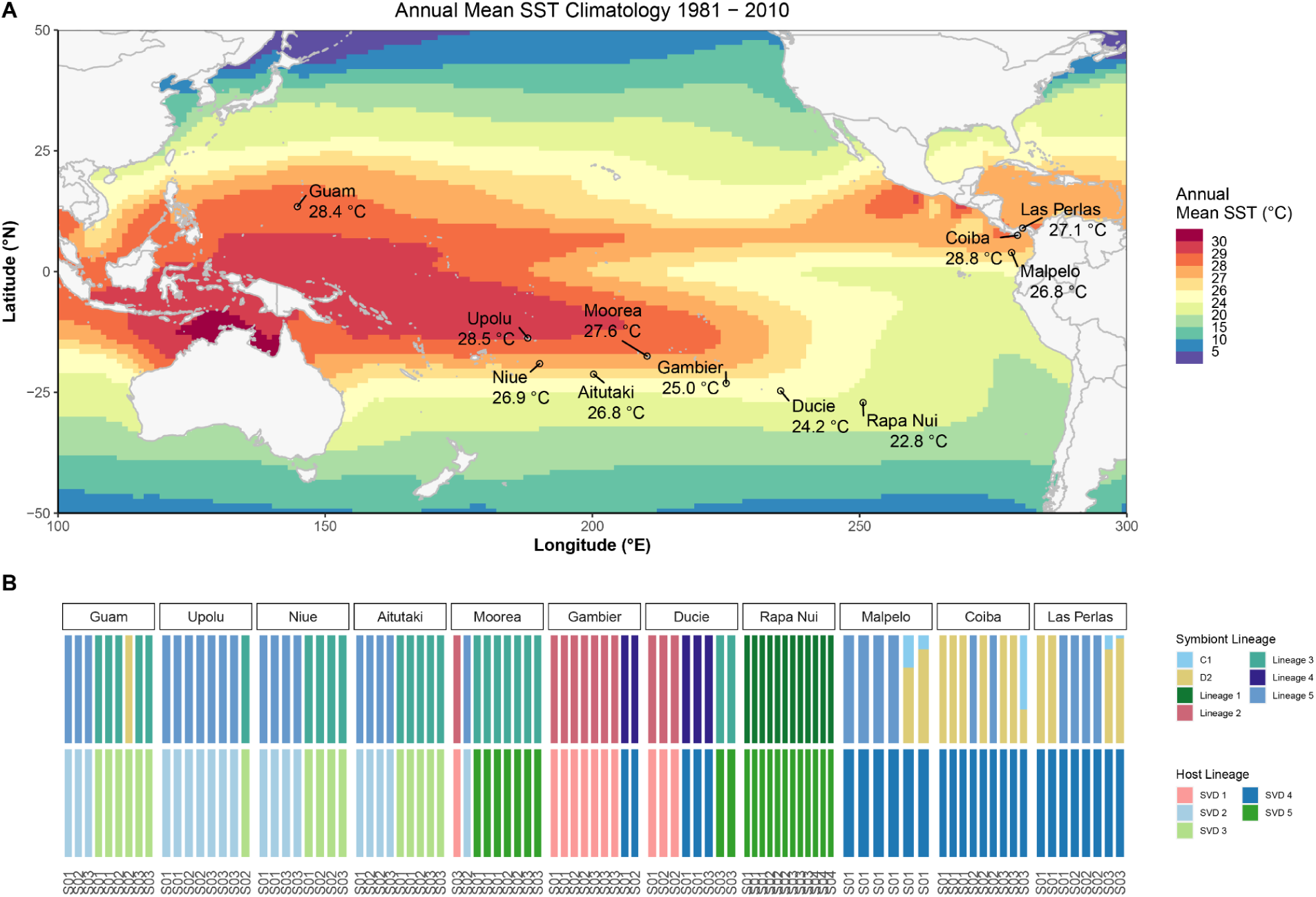
Pocillopora and Symbiodiniaceae lineages identified in each sample across the Pacific Ocean. (A) Map showing the 11 islands sampled during the *Tara* Pacific expedition. (B) Lineage assignations of sampled colonies for both the dominant *Cladocopium C1* genetic lineage (top, including proportions of *Durusdinium D2* when it was also present) and *Pocillopora* host genotype (middle). Also displayed are historical mean sea surface temperatures (SST) at each sampling site (bottom). Lineages were identified from an analysis of single nucleotide polymorphisms (SNPs) performed using metagenomic reads for the *Pocillopora* host and on metatranscriptomic reads for the Symbiodiniaceae.

Using the ITS2 metabarcoding data processed with the SymPortal method (see paper number 6), we identified a total of 28 ITS2 profiles across *Pocillopora* colonies (Figure S2). Whereas the majority of colonies (84 in total) displayed association with *Cladocopium* symbionts, we identified 9 colonies containing only *Durusdinium (*D2 *ITS2 sequence)* and 9 colonies containing both *Cladocopium* (mainly C42a ITS2 sequence) and *Durusdinium* (D2 ITS2 sequence) in varying proportions. *Durusdinium*-containing colonies were restricted to *Pocillopora* SVD4 hosts in the Eastern Pacific (Las Perlas, Coiba, and Malpelo) and one sample in *Pocillopora* SVD3 on Guam (Figure 1B).

Clustering of these *Cladocopium* populations into robust lineages based on the ITS2 was not possible due to the high diversity and the intragenomic variability of the ITS2 sequences. Therefore, we performed a single nucleotide polymorphism (SNP) analysis on the 82 colonies containing only a single *Cladocopium* species to identify major photosymbiont lineages present in the *Pocillopora* samples. SNPs were called by aligning the metatranscriptomic reads on the coding sequences of the *Cladocopium goreaui* genome resulting in the detection of 3,712 SNPs distributed across 1,354 transcripts within the 82 colonies (see methods). The hierarchical clustering of these SNPs revealed five lineages of *Cladocopium*, hereafter referred to as L1 to L5 (Figure 1B and Figure S3). We then compared the ITS2 profile distances between each pair of *Cladocopium*-containing *Pocillopora* colonies (see methods section 2.3.2.) to the SNP-based clustering of *Cladocopium* populations (Figure S4). In general, we observed a high degree of overlap between the two identification methods across *Cladocopium* lineages. However, whereas L1 and L5 are clearly isolated based on the ITS2 profile distances, L2, L3 and L4 are more similar to one another in the principal coordinate analysis and would have been difficult to identify using only ITS2 profiles. The ITS2 profile distances indicate that *Cladocopium* lineages which were undefined in the SNP clustering method belong to L5.

Lineage 1 is genetically distant from all other lineages and restricted to Rapa Nui island. L2 and L4 are both restricted to the central Pacific (Ducie, Gambier and Moorea islands). L2 is specific to the SVD1 lineage of the *Pocillopora* host and L4 to *Pocillopora* SVD4. Finally, L3 and L5 are both present in the Western Pacific (Aitutaki, Niue, Upolu, and Guam) and are also host specific in this oceanic region (L3 in *Pocillopora* SVD3 and L5 in *Pocillopora* SVD2; Figure 1B). Interestingly, L5 is also present in SVD4 in the East Pacific. In general, we observed high host-symbiont specificity at basin-wide scale in all *Pocillopora* lineages except within *Pocillopora* SVD4, which is in symbiosis with *Cladocopium C1* and *Durusdinium,* and *Pocillopora* SVD5 which hosted both *Cladocopium* L1 and L3 symbionts.

### 3.2. Identifying the main drivers of holobiont gene expression and functional divergences

#### 3.2.1. Variation partitioning analysis

The variation partitioning approach allowed us to identify genes whose expression levels were driven primarily by the sampling environment, the host genetic lineage, or the algal genotype (see methods). For the host, among the 1,821 genes with greater than 50% of their expression variation attributed to these three factors, 57% (1,042 genes) are linked to the environment, 33% (606 genes) are associated with the host genetic lineage, and 9.5% (173 genes) are attributed to the symbiont genetic lineage (Figure 2A). For the *Cladocopium C1* photosymbiont, we identified 2,424 genes above the 50% explained variation threshold, with 80% of the genes (1,950 genes) best explained by the genetic lineage of the symbiont, and only 10% (251 genes) and 9% (223 genes) of the genes linked to the sampling island and the genetic clade of the *Pocillopora* host, respectively (Figure 2B).

**Figure 2.**
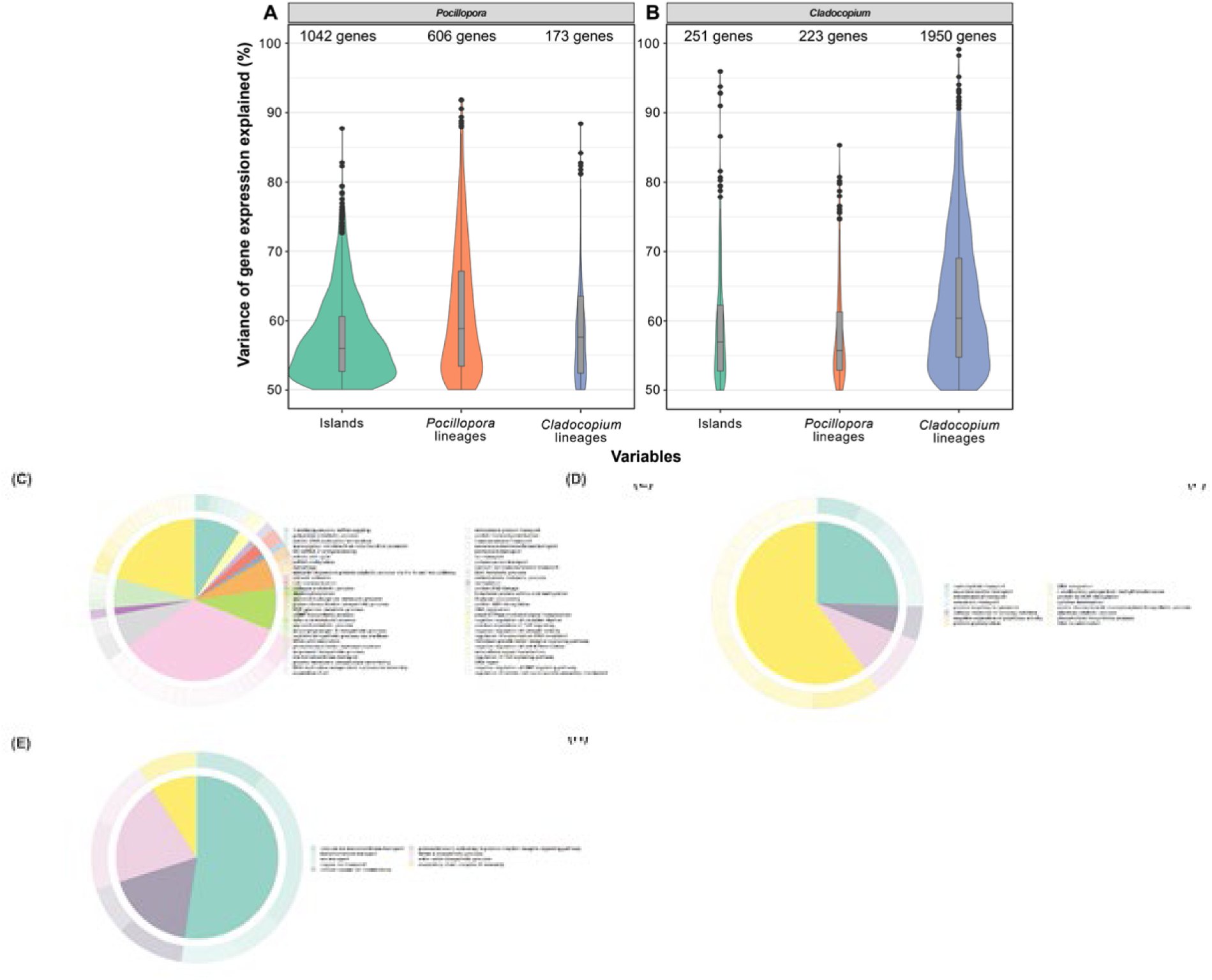
Contribution of environmental and genetic variables to Pocillopora and Cladocopium gene expression and biological process functional enrichments of top variant genes. Distribution of genes in which more than 50% of the variation in their expression is explained by one of three predictor variables: the sampling island, the *Pocillopora* genetic lineage, and the *Cladocopium* lineage. The number of genes in each distribution is indicated above each violin plot. A) Gene expression of *Pocillopora* across 101 samples. B) Gene expression of *Cladocopium* across 70 samples. Samples of Rapa Nui island and those containing *Durusdinium* symbiont were excluded. Host functional enrichments among top variant genes are shown for those best explained by C) island of sampling, D) the host genetic lineage, and E) the photosymbiont genetic lineage.

We investigated functional enrichments among these top variant genes and the results are presented in Tables S1-S6. In the *Pocillopora* host, genes with expression dependent on the island of sampling were enriched in biological process ontologies related to lipid and carbohydrate metabolism (and 17/125 genes, respectively), ion transmembrane transport (16/82 genes) including transmembrane transport of ammonium (3/11 genes) and calcium (9/64 genes), as well as regulation of DNA repair (9/102 genes) and autophagy (3/11 genes; Table S1, Figure 2C). In Cladocopium, genes whose expression is dependent on the island of sampling were enriched in PFam domains associated with heat shock chaperones (HSP70, 2 genes), active transport of protons across membranes (E1-E2 ATPases, 3 genes), and oligosaccharide processing (Glucosidase II beta subunit-like, 2 genes; Table S2).

Host genes whose expression was dependent on the Pocillopora genetic lineage were enriched in biological processes related to carbohydrate (3/29 genes) and xenobiotic transport (3/77 genes), DNA recombination (8/156 genes), and response to ionizing radiation (1/1 gene; Table S3, Figure 2E). *Cladocopium C1* genes whose expression was dependent on the photosymbiont lineage included PFam enrichments related to ion transport (57 genes) and papain family cysteine proteases (10 genes) which are known to show increased expression in plants in response to multiple environmental stressors (Liu et al. 2018a; Table S4).

*Pocillopora* genes whose expression was dependent on the photosymbiont lineage were enriched in biological process ontologies related to ion transmembrane transport (6/256 genes) particularly of calcium (3/64 genes), nitric oxide biosynthesis (1/1 gene), and assembly of the mitochondrial respiratory chain complex IV (1/2 genes; Table S5). Among *Cladocopium* genes whose expression is dependent on the host, 14 are involved in carbohydrate metabolism (Galactose oxidase, Glucan synthase, Glycosyl hydrolase) and gluconeogenesis (Fructose-bisphosphate aldolase, Enolase, Phosphofructokinase-2). Thirteen calcium-transporter and calcium-dependant protein kinases that have been shown to play a role in the establishment of symbiosis (Rosic *et al*. 2015) are also dependent on the host lineage (Table S6).

#### 3.2.2. Discriminant analysis of principal components (DAPC)

To determine the main factors controlling the gene expression variations in the host and the symbiont, a discriminant analysis of principal components (DAPC) was performed. DAPC is a multivariate analysis which transforms the data using a principal component analysis (PCA) then identifies clusters using a discriminant analysis. The number of principal coordinates (PCs) and discriminant functions (DF) retained, the proportion of colonies correctly reassigned to their *a priori* groups, and the overall proportion of gene expression variance explained were calculated for the island of sampling, the primary genetic lineage, and the genetic lineage of the symbiotic partner using the gene expression of the host or the gene expression of the photosymbiont (Table S9).

For the host, the 12 PCs retained after a-score maximization in the DAPC explained 56.68% of total gene expression variance (Figure S5A). Discriminant axis eigenvalue scores were roughly equivalent between DF1 and DF2 for all three grouping models (Figure S5B-D). For the photosymbiont, the 11 PCs retained explained 73.86% of total gene expression variance (Figure S5E). Discriminant axis eigenvalue scores were highest for DF1 for all three grouping models (Figure S5F-H).

Based on proportions of colonies correctly reassigned to pre-defined groups, DAPC revealed marginally stronger influence of genotype (both primary lineage and of the symbiotic partner) than of the environment for both the host and the symbiont. When colonies were assigned to pre-defined groups based on the environment (island of sampling) we were able to correctly reassign 96% and 91% of colonies based on *Pocillopora* and *Cladocopium* expression profiles, respectively (Table S9, Figure 3A and B). Unconstrained analyses also confirmed that the distinction between environments of sampling was significant for both the host and photosymbiont (Table S10, PERMANOVA p < 0.001).

**Figure 3.**
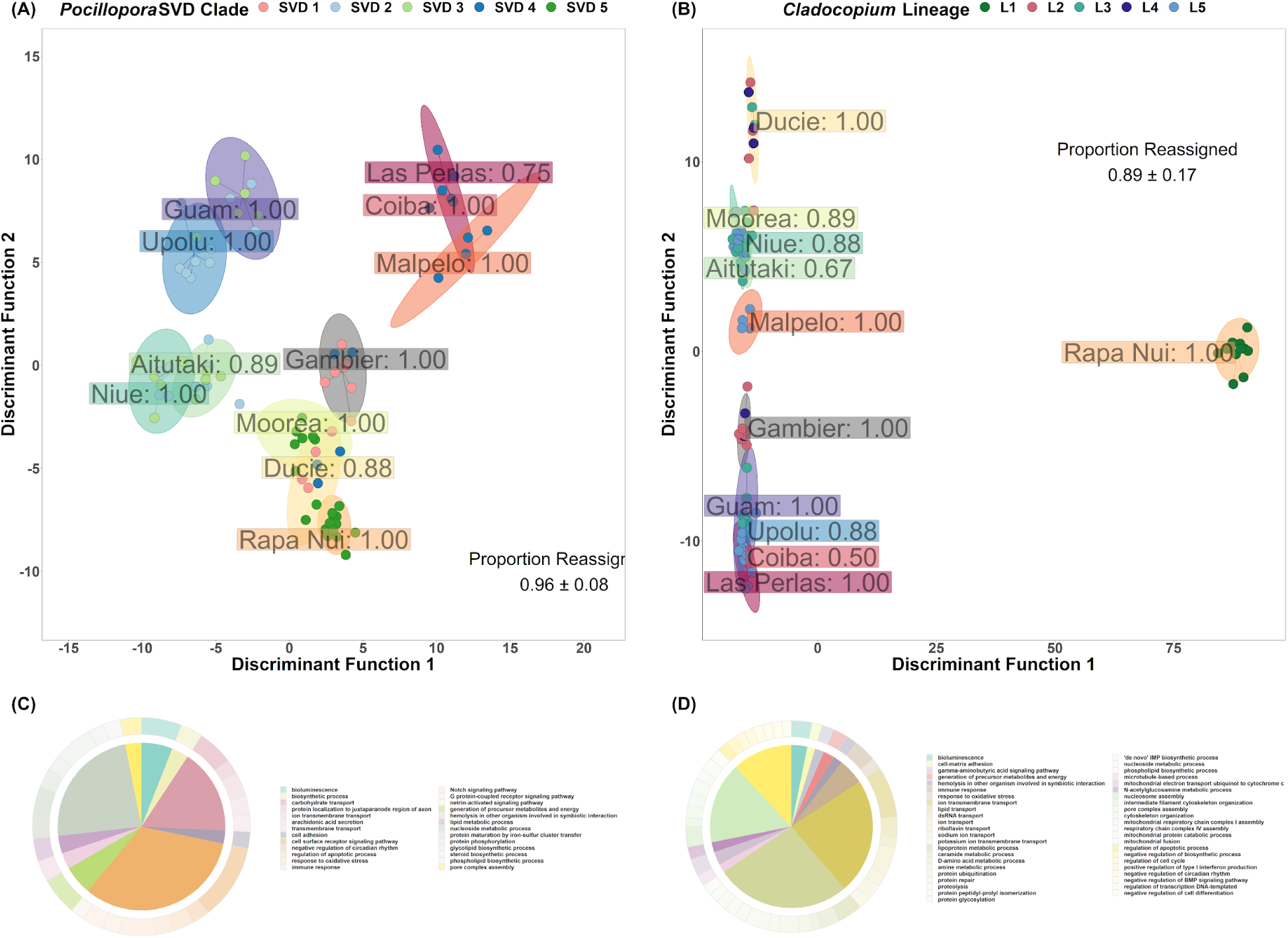
Discriminant analysis of principal components (DAPC) of gene expression data grouped by environment and biological process functional enrichments of discriminant genes in the Pocillopora host (left) and Cladocopium photosymbiont (right). DAPC scatter plots showing expression profiles for the host (A) and algal symbiont (B) when colonies were grouped by sampling island. Points represent individual colony expression profiles and are colored by genetic lineage. Shaded ellipses denote 95%-confidence intervals around the group (island) mean. Group-specific proportions of correct reassignments are indicated within each cluster (boxed labels) and overall model proportions of correct reassignment (mean ± standard deviation) are presented within each panel. Also shown are functional enrichments among host genes contributing to the two discriminant functions in the DAPC, DF1 and DF2, (C and D, respectively).

When colonies were assigned to pre-defined groups based on primary genetic lineage, proportions of correct reassignment were maximized at 100% for both the host and algal symbiont (Table S9, Figure 4A and B). These genetic distinctions were also confirmed to be significant in unconstrained analyses for both the host and algal symbiont (Table S10, PERMANOVA p < 0.001). Similarly, because of the strong host-symbiont fidelity we observed, when colonies were grouped by the genetic lineage of their symbiotic partner, we were able to correctly reassign 99% of colonies in the host and 100% of colonies in the algal symbiont (Table S9, Figure 5A and B). These partner distinctions were also significant in unconstrained analyses for both the host and algal symbiont (Table S10, PERMANOVA p < 0.001). These results indicate that the main driver of gene expression is the genetic lineage for both the host and photosymbiont with geography (i.e., local environment) being secondary.

**Figure 4.**
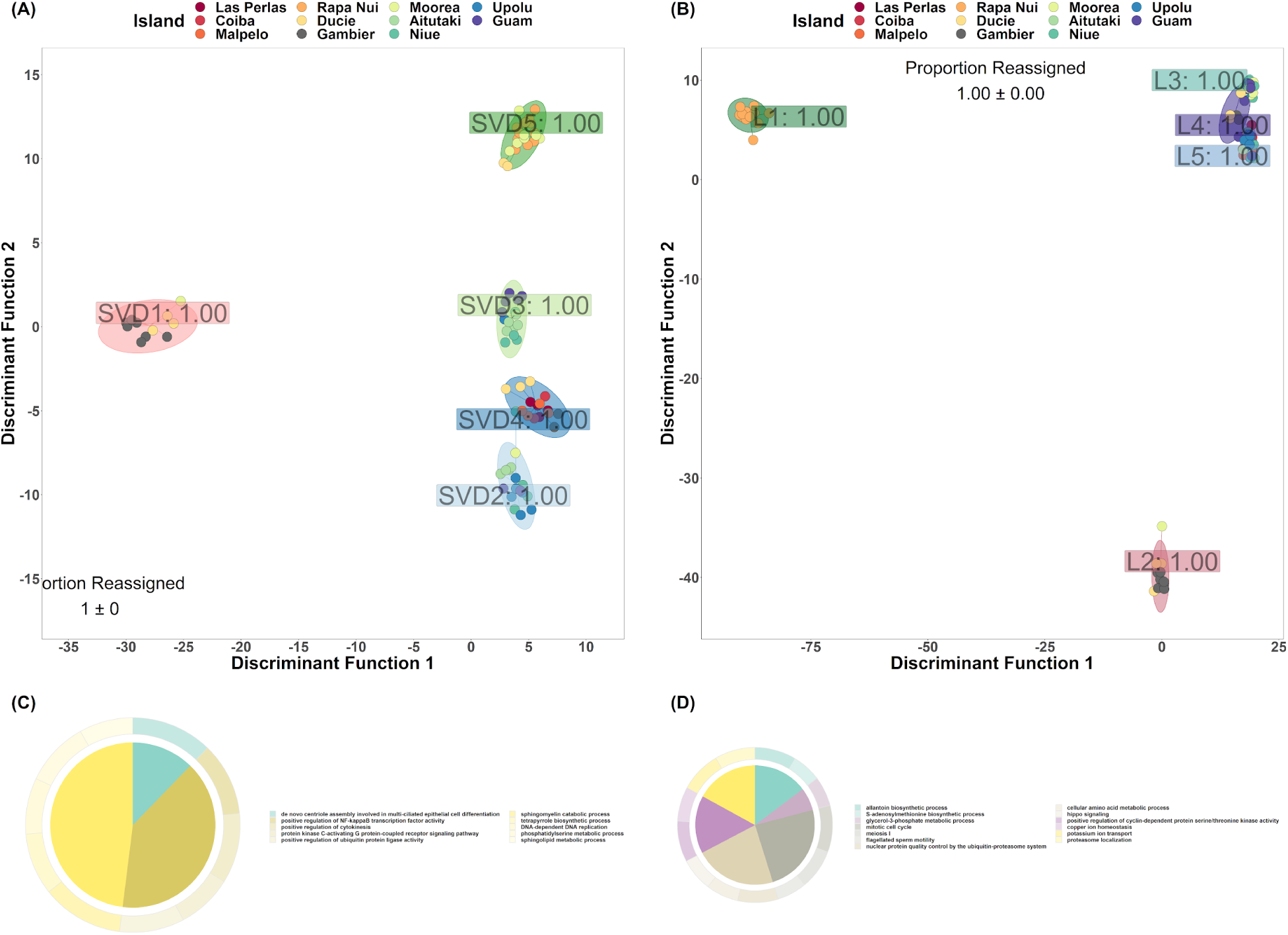
Discriminant analysis of principal components (DAPC) of gene expression data grouped by primary genetic lineage and biological process functional enrichments of discriminant genes in the Pocillopora host (left) and Cladocopium photosymbiont (right). DAPC scatter plots showing expression profiles for the host (A) and algal symbiont (B) when colonies were grouped by primary genetic lineage. Points represent individual colony expression profiles and are colored by island of sampling. Shaded ellipses denote 95%- confidence intervals around the group (lineage) mean. Group-specific proportions of correct reassignments are indicated within each cluster (boxed labels) and overall model proportions of correct reassignment (mean ± standard deviation) are presented within each panel. Also shown are functional enrichments among host genes contributing to the two discriminant functions in the DAPC, DF1 and DF2, (C and D, respectively).

**Figure 5.**
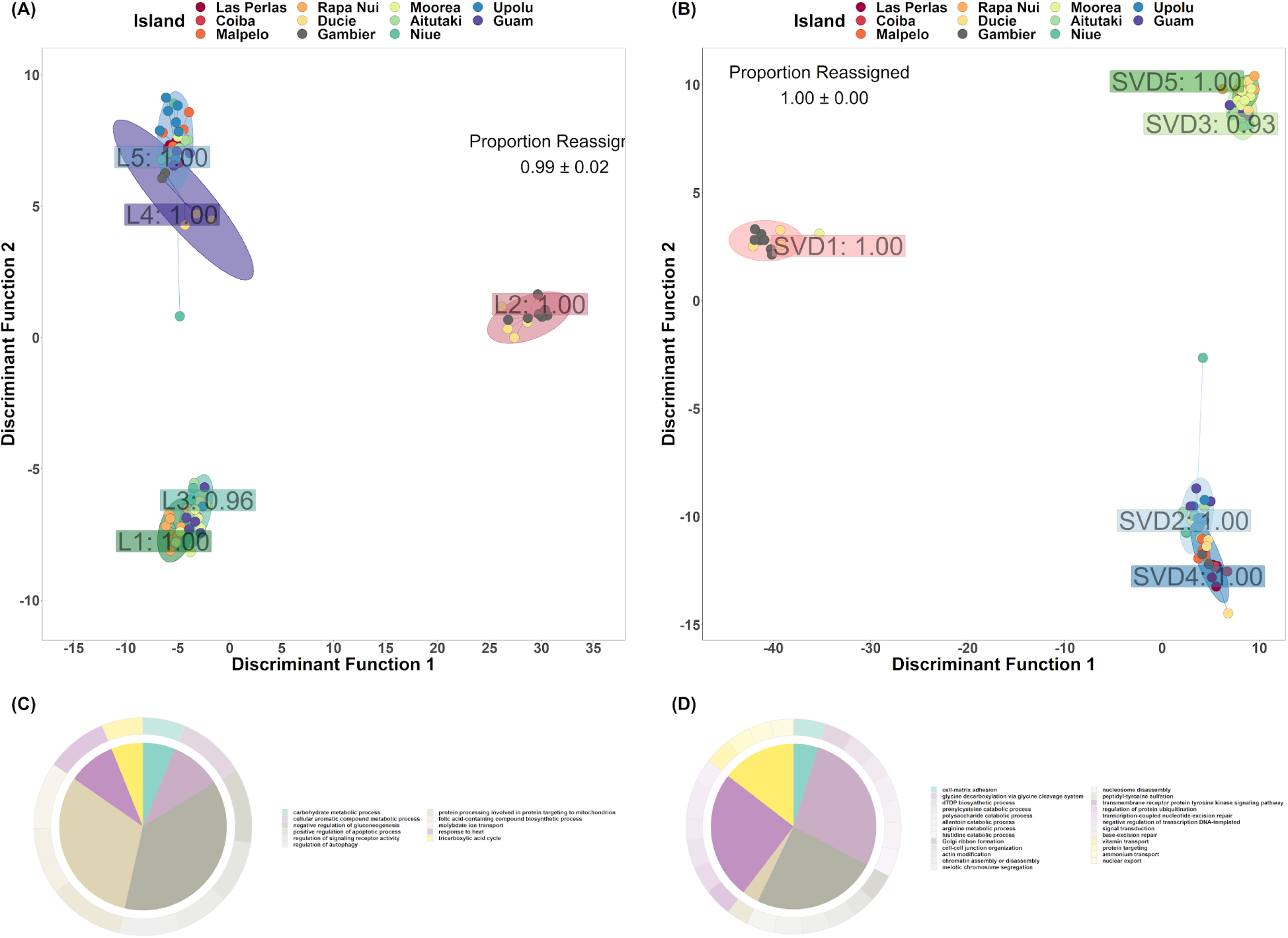
Discriminant analysis of principal components (DAPC) of gene expression data grouped by symbiotic partner and biological process functional enrichments of discriminant genes in the Pocillopora host (left) and Cladocopium photosymbiont (right). DAPC scatter plots showing expression profiles for the host (A) and algal symbiont (B) when colonies were grouped by genetic lineage of the symbiotic partner. Points represent individual colony expression profiles and are colored by island of sampling. Shaded ellipses denote 95%- confidence intervals around the group (symbiotic lineage) mean. Group-specific proportions of correct reassignments are indicated within each cluster (boxed labels) and overall model proportions of correct reassignment (mean ± standard deviation) are presented within each panel. Also shown are functional enrichments among host genes contributing to the two discriminant functions in the DAPC, DF1 and DF2, (C and D, respectively).

In the coral host under the environmental model, DAPC differentiated expression profiles by geography along DF1 and by environmental factors (especially historical SST) along DF2. Colonies from the Eastern Pacific (Isla de Las Perlas, Coiba, and Malpelo) clustered around positive values of DF1 whereas colonies from the Western Pacific (Aitutaki, Niue, Upolu, and Guam) clustered at the negative end of this axis. Along DF2, colonies from islands with historically elevated SST values (Upolu, Guam, Isla de Las Perlas, Coiba, and Malpelo) clustered around positive DF2 values and colonies from the coldest islands (Rapa Nui and Ducie) clustered around negative values (Figure 3A). In the *Cladocopium* photosymbiont under the environmental model, DF1 primarily separated colonies from Rapa Nui from all others. The DF2 axis followed a similar pattern to that observed in the host, namely that colonies were distinguished from one another on this axis based on historical SST of the island of sampling (Figure 3B).

Under the host primary genetic model, DF1 primarily distinguished *Pocillopora* SVD 1 colonies from all others. The DF2 axis served to separate the remaining four SVD clades from one another, with clades SVD 2 and 4 (sister groups) clustered near one another at negative DF2 values, clades SVD 3 and SVD 1 overlapping around DF2 equal to zero, and SVD5 clustering at positive DF2 values (Figure 4A). In *Cladocopium*, DF1 of the primary genetic model, distinguished Lineages 1 (the most divergent lineage) and 2 from all others. DF2 served primarily to distinguish Lineage 2 from all others(Figure 4B).

Finally, under the symbiotic partner model, host colonies containing *Cladocopium* L2 symbionts were strongly separated from all others along DF1 whereas DF2 primarily distinguished colonies containing L1 and L3 photosymbionts (negative DF2 values) from those containing L4 and L5 photosymbionts (positive DF2 values, Figure 5A). In *Cladocopium*, DF1 of the symbiotic partner model distinguished expression profiles from photosymbionts inhabiting SVD 1 corals from all others and DF2 distinguished between photosymbiont inhabiting SVD 2 and SVD 4 colonies (negative DF2 values) from those inhabiting SVD 3 and SVD 5 colonies (positive DF2 Values, Figure 5B).

We also investigated gene ontology enrichments among genes which contributed most strongly to the two discriminant functions in each model (i.e., discriminant genes). The results of these analyses are presented in Tables S1-S6. Under the environmental model, host genes which discriminated between islands along DF1, the “geographical axis”, were enriched in biological processes related to carbohydrate binding and transport (27 and 15 genes, respectively) and glycolipid biosynthesis (13 genes, Figure 3C). Host genes which discriminated along DF2, the “SST axis”, were enriched in processes related to protein ubiquitination and repair (32 and 2 genes, respectively) and mitochondrial respiratory chain complex assembly (4 genes, Figure 3D). Functional enrichments shared across both axes included processes related to ion transmembrane transport (28/45 genes for DF1/DF2, respectively), generation of precursor metabolites and energy (2/1/4 genes for DF1/DF2/shared, respectively), immune response (3/4/7 genes), response to oxidative stress(2/3/8 genes), and cytolysis in other organism involved in symbiotic interaction (3 shared genes, Figure 3C and E). In *Cladocopium*, genes whose expression strongly distinguished photosymbionts from Rapa Nui from all others (DF1 contributing genes) were enriched in protein family (Pfam) domains related to chlorophyll A-B binding (35 genes) and ion transport (91 genes; Table S2). *Cladocopium* genes which contributed to DF2 were not significantly enriched in any Pfam domains (Table S2).

Under the primary genetic lineage model, DF1 identified *Pocillopora* genes which distinguished SVD 1 colonies from all other lineages. These genes were enriched in biological processes related to carbohydrate transport (14 genes), lipid biosynthesis (5 genes), immune response (8 genes), tumor necrosis factor-mediated signaling (3 genes), and regulation of response to virus (3 genes, Figure 4C). Host genes which contributed most strongly to expression profile discrimination along DF2 were enriched in processes related to cytolysis in other organism involved in symbiotic interaction (3 genes), ion transport (90 genes), and protein glycosylation and phosphorylation (26 and 197 genes, respectively, Figure 4D). Functional enrichments shared across both axes included processes related to calcium ion and xenobiotic transport (9/12/16 and 14/16/15 DF1/DF2/shared genes, respectively), lipid biosynthesis and transport (1/1/4 and 6/2/8 genes, respectively), and regulation of apoptosis (28/31/16 genes, Figures 4C and E). Host genes which were unique to the primary genetic lineage model (i.e., not shared with the symbiotic partner model) were principally involved in cytokinesis cell division. These genes were enriched in processes related to centriole replication (1 gene), positive regulation of cytokinesis (1 gene), DNA replication (1 gene), and sphingolipid metabolism (1 gene) along DF1 and processes related to mitosis and meiosis (2 and 1 genes, respectively) and glycerol−3−phosphate metabolism (1 gene) along DF2. In *Cladocopium*, DF1 of the primary lineage model was enriched in PFam domains related to chlorophyll A-B binding (37 genes) and ion transport (104 genes; Table S4). *Cladocopium* genes which contributed to DF2 were enriched in Pfam domains related to ion transport (113 genes) and P-loop containing dynein motor region (13 genes, Table S4).

According to the symbiotic partner model, host genes which distinguished *Cladocopium* L2-containing colonies (DF1 genes) from all others were enriched in biological processes related to carbohydrate biosynthesis, binding, and transport (8, 15, and 28 genes, respectively), mitochondrial cytochrome c oxidase assembly (3 genes), tumor necrosis factor-mediated signaling (3 genes), and regulation of response to virus (3 genes, Figure 5C). Host genes which discriminated between colonies along DF2 were enriched in processes related to cytolysis in another organism involved in symbiotic interaction (4 genes), phospholipid biosynthesis (8 genes), and nucleosome assembly (8 genes, Figure 5D). Functional enrichments shared across both axes included processes related to xenobiotic and calcium ion transmembrane transport (11/20/14 and 12/10/16 DF1/DF2/shared genes, respectively), carbohydrate transport (4/5/10 genes), immune response (4/2/5 genes), lipid biosynthesis (1/1/4 genes), regulation of apoptosis (26/28/19 genes). Genes unique to this model and which contributed strongly to DF1 were enriched in processes related to the TCA cycle (1 gene), negative regulation of gluconeogenesis (1 genes), positive regulation of apoptosis (1 genes), regulation of autophagy (1 genes), and response to heat (1 genes). Unique genes contributing to DF2 were enriched in processes related to polysaccharide catabolism (1 genes), meiotic chromosome segregation and chromatin assembly/disassembly (1 gene each), base-excision repair (1 gene), and ammonium transport (2 genes). In *Cladocopium*, DF1 of the DAPC discriminated expression profiles of photosymbionts residing in SVD1 hosts from all others and genes contributing to this axis were enriched in Pfam domains related to chlorophyll A-B binding (37 genes) and ion transport (104 genes; Table S6). Genes which contributed to separation of colonies along DF2 were enriched in Pfam domains related to ion transport (113 genes), and P-loop containing dynein motor regions (13 genes; Table S6).

#### 3.2.3. Expression variance and evolution model (EVE) diverse and divergent genes

In the *Pocillopora* host, we identified a total of 8,378 potential divergent candidate genes. There were no diverse candidate genes identified in the coral host. Host genes whose expression profiles diverged significantly between SVD clades were enriched in biological processes related to G-protein coupled receptor signaling (692/2139 genes) including activation of the innate immune response (5/9 genes), ATPase-coupled transmembrane transporter activity (47/108 genes) including transport of calcium (28/64 genes) and xenobiotic compounds (35/77 genes), carbohydrate transport (13/29 genes), glycolipid biosynthesis (11/25 genes), and cytolysis of another organism involved in a symbiotic interaction (3/4 genes).

In the *Cladocopium C1* photosymbiont, we identified 146 and 17,846 potential diverse and divergent gene candidates, respectively. We observed no Pfam enrichments among photosymbiont genes displaying diverse within-lineage expression. However, *Cladocopium C1* genes with expression profiles that diverged significantly between lineages were enriched in Pfam domains related to ion transport (145 genes), chlorophyll A-B binding (39 genes), and cyclic nucleotide-binding (39 genes).

#### 3.2.4. Constrained correspondence analysis

Using a constrained correspondence analysis (CCA), we identified nine environmental variables and one genetic variable from among those that were not strongly autocorrelated that best explained the dispersion among host gene expression profiles (Figure 6A, Table S11). Similarly, for the photosymbiont, we identified eight environmental variables and one genetic variable that best explained the dispersion among gene expression profiles (Figure 6B, Table S11).

**Figure 6.**
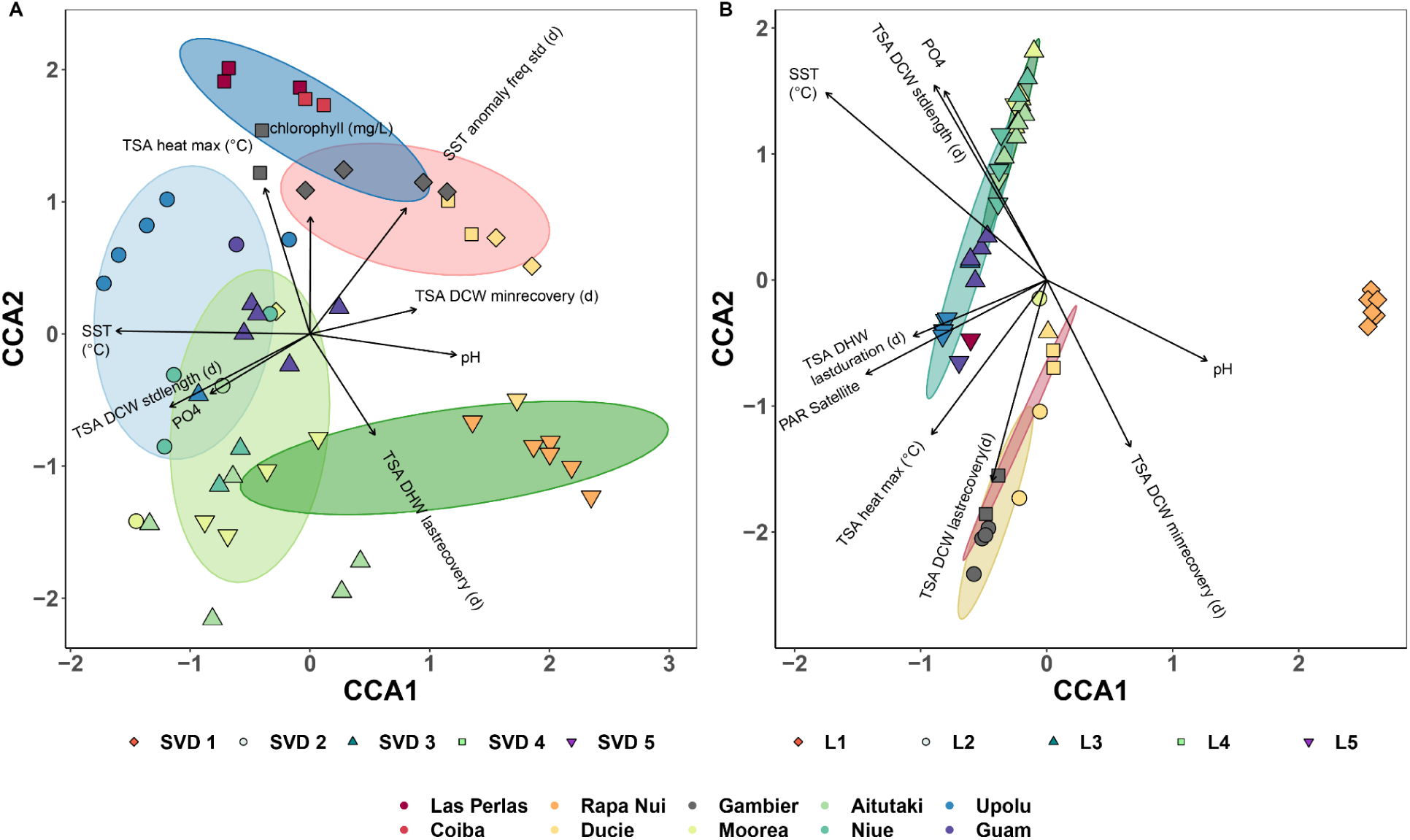
Constrained correspondence analysis (CCA) of Pocillopora spp. host and Cladocopium C1 symbiont gene expression profiles from colonies collected across the Pacific Ocean. Scatter plots show (A) Pocillopora host and (B) Cladocopium C1 photosymbiont expression profiles for individual colonies with data colored by island of collection and clustered by genetic lineage (i.e., Pocillopora SVD clade and Cladocopium C1 genetic lineage, respectively; shaded ellipses). Length (strength) and direction (towards increasing values) of vector arrows denote the relative influence of the top explanatory environmental variables (permuted p-value ≤ 0.01) for each dataset.

For both the coral host and *Cladocopium C1* photosymbiont, mean climatological SST and primary genetic lineage were the two strongest contributors to gene expression variation between colonies. In the Pocillopra host, separation of gene expression profiles along constrained correspondence axis 1 (CCA1 Figure 6A) was primarily driven by temperature and pH. Colonies from islands with high pH values (e.g., Rapa Nui) clustered around positive CCA1 values whereas colonies from islands with historically elevated mean SST (e.g., Upolu) clustered around negative CCA1 values. In contrast, CCA2 was primarily tied to SST variability (Figure 6A). Positive CCA2 values were associated with islands with the highest maximum surface temperature anomalies (e.g., Las Perlas and Cobia) or those with large standard deviations in the frequency of elevated SST anomaly events (e.g., Gambier). Additionally, positive CCA2 values were strongly correlated with elevated environmental chlorophyll concentrations suggesting a link to potential eutrophication. Conversely, negative CCA2 values were associated with islands with long recovery periods between extreme heating events (i.e., large degree heating week recovery period, for example Rapa Nui) and/or long degree cooling week durations (e.g., Moorea).

In the *Cladocopium C1* photosymbiont, no single environmental variable was strongly correlated with CCA1, although pH and photosynthetically active radiation (PAR) both contributed somewhat to gene expression separation along this axis (Figure 6B). Again, colonies from islands with high pH values (e.g., Rapa Nui) clustered around positive CCA1 values whereas those closer to the equator, and therefore with higher incident PAR, tended to cluster around negative CCA2 values (e.g., Upolu). As in the coral host, CCA2 was primarily associated with temperature variability, however the specific associations differed. For example, exposure to extreme cooling events played a more significant role in structuring photosymbiont expression profiles than in the coral host (Figure 6B). Colonies from islands with long recovery periods between cooling events (i.e., Gambier and Ducie) tended to cluster around negative CCA2 values. Conversely, colonies from islands with prolonged exposure to cooling events or with elevated SSTs and environmental phosphate concentrations (e.g., Aitutaki and Niue) clustered together around positive CCA2 values. These results suggest that whereas gene expression profiles in both the coral host and photosymbiont are primarily structured by climatological SST, it is the frequency of exposure to extreme heating/cooling events that drives substructuring of gene expression in the host/photosymbiont, respectively.

### 3.3. Differential gene expression following symbiont shuffling

According to unconstrained analysis, global gene expression profiles of colonies from the Eastern Pacific (Isla de Las Perlas, Coiba, and Malpelo) differed between islands (PERMANOVA, p ≤ 0.05), but not between *Cladocopium C1*- and *Durusdinium D2*-containing colonies (Figure S6, PERMANOVA p > 0.05). In addition, intra-island comparison of expression profiles between colonies in the Eastern Pacific revealed only a small number of genes that were differentially expressed between *C*- and *D*-containing hosts. Because the DEG counts were dependent on the number of colonies sampled from each location, we also calculated colony-normalized averages using a repeated random subsampling approach. We observed 294 ± 363, 129 ± 53, and 110 ± 15 genes up-regulated in D-containing colonies (p ≤ 0.001 and a LFC ≥ 2) on the three islands, respectively (Figure S7A). We also observed 171 ± 164, 211 ± 102, and 200 ± 59 genes down-regulated in D-containing colonies (p ≤ 0.001 and a LFC ≤ 2) on the three islands, respectively (Figure S7B). Very few DEGs were shared between pairs of islands. The number of pairwise shared DEGs ranged from 3 (up-regulated in both Las Perlas and Coiba) to 9 (down-regulated in both Las Perlas and Coiba) and no gene was commonly differentially expressed across all three islands (Figure S7A and B).

We also examined functional enrichments among genes differentially expressed between *C*- and D-containing colonies and these results are listed in Table S2. In general, genes up-regulated in D-containing colonies on Las Perlas, were enriched in GO biological processes related to bioluminescence (GO:0008218), xenobiotic transport (GO:0042908), gluconeogenesis (GO:0006094), mitochondrial protein catabolic process (GO:0035694), base-excision repair (GO:0006284), and transmembrane transport (GO:0055085; Table S2). Genes down-regulated in D-containing colonies on Las Perlas, were enriched in GO biological processes related to calcium ion transmembrane transport (GO:0070588), and phospholipid biosynthetic processes (GO:0008654). Genes up-regulated in D-containing colonies on Coiba, were enriched in GO biological processes related to protein phosphorylation (GO:0006468) and proteolysis (GO:0006508; Table S2) whereas down-regulated genes were enriched in processes related to chaperone-mediated protein folding (GO:0061077), chaperone-mediated protein transport (GO:0072321), synaptic vesicle transport (GO:0048489), nuclear envelope organization (GO:0006998), protein phosphorylation (GO:0006468), cell adhesion (GO:0007155), neuron projection development (GO:0031175), and intermediate filament cytoskeleton organization (GO:0045104). Finally, genes up-regulated in D-containing colonies on Malpelo, were enriched in GO biological processes related to bioluminescence (GO:0008218), regulation of signaling receptor activity (GO:0010469), vacuolar transport (GO:0007034), and xenobiotic transport (GO:0042908; Table S2). Genes down-regulated in D-containing colonies on Malpelo, were enriched in GO biological processes related to bioluminescence (GO:0008218), regulation of apoptotic process (GO:0042981), protein phosphorylation (GO:0006468), S-adenosylmethionine biosynthesis (GO:0006556), regulation of pH (GO:0006885), notch signaling pathway (GO:0007219), MyD88-dependent toll-like receptor signaling pathway (GO:0002755), proteolysis (GO:0006508), signal transduction (GO:0007165), protein ubiquitination (GO:0016567), and netrin-activated signaling pathway (GO:0038007).

## 4. DISCUSSION

In this study, we identify and discuss the high host-photosymbiont fidelity we observed among Pocillopora lineages across environments. We also investigate instances of potential symbiotic flexibility and/or breakdown which we observed among corals living on reefs which have historically experienced elevated sea surface temperatures (SSTs). We propose that these mechanisms represent a three-tiered strategy of thermal acclimatization/adaptation in *Pocillopora* corals and discuss the implications of this strategy for continued persistence of these holobionts under future ocean warming.

### 4.1. Pocillopora corals exhibit high symbiotic fidelity with distinct Cladocopium lineages

Within *Pocillopora*, it is well known that a large diversity of Symbiodiniaceae species are found in symbiosis with *Pocillopora* colonies (LaJeunesse et al. 2007, 2010; Stat et al. 2009; Cunning et al. 2013). In total, we observed five lineages of *Cladocopium C1* and one strain of *Durusdinium* (D2) in symbiotic association with *Pocillopora* host colonies across the Pacific ocean (Figure 1B). Of the five *Pocillopora* lineages that we sampled, three exhibited near perfect symbiotic fidelity with distinct lineages of Cladocopium (i.e., one photosymbiont lineage for each host; Figure 1B) and this fidelity persisted regardless of the environment of sampling. For example, Pocillopora SVD 1 colonies, present on Ducie, Gambier, and Moorea, were always found in symbiosis with *Cladocopium C1* L2 symbionts. Similarly, Pocillopora SVD 2 colonies, present on several islands in the Western Pacific, always hosted *Cladocopium C1* L5 as their primary photosymbiont. Colonies of Pocillopora SVD 3 were present in both the Central and Western Pacific and were, save for one instance which we discuss in section 4.5 below, always found in association with *Cladocopium C1* L3. Strong fidelity between a single coral host and a single Symbiodiniaceae lineage has been widely observed across a variety of scleractinian species (Pinzón and LaJeunesse 2011; Baums et al. 2014; Thornhill et al. 2014; Howells et al. 2020). Our findings therefore support the general consensus that fidelity to a single clade of symbiont is the dominant pattern in most scleractinian corals (Baird et al. 2007).

Despite this overarching tendency towards symbiotic fidelity, we also detected two broad instances of imperfect specificity (i.e., symbiont flexibility) which warrant further discussion. In the first case, we observed a switch in symbiotic association between colonies of the same host lineage inhabiting different environments. For example, Pocillopora SVD 4 colonies present on Rapa Nui (mean climatological SST 22.8 °C) were found to be in symbiosis with *Cladocopium C1* L1 symbionts. However, when present on Ducie or Moorea (climatological SSTs of 24.2 °C and 27.6 °C, respectively), this same host lineage was found to be associated with *Cladocopium C1* L3 symbionts (Figure 1B). Similarly, host SVD 4 colonies present in the Eastern Pacific (mean climatological SST of 27.6 ± 1.1 °C for the three islands) were primarily in symbiosis with *Cladocopium C1* L5 (although *Durusdinium D2* was also present, see section 4.5, below). However, SVD 4 colonies from the Central Pacific (mean SST of 24.6 ± 0.5 °C for the two islands) contained only *Cladocopium C1* L4 photosymbionts. This observation suggests that both these coral hosts display flexibility in their symbiotic associations with Cladocopium across their range and that this flexibility may be dependent on the local thermal environment (Berkelmans and van Oppen 2006; Abrego et al. 2009). In both instances, photosymbionts belonging to lineages L3 and L5 seem to have been preferentially selected for by *Pocillopora* hosts when present on islands with historically elevated SSTs. In the second broad case of symbiotic flexibility, we observed a switch from *Cladocopium C1* to *Durusdinium D2* as the dominant photosymbiont in two *Pocillopora* lineages. This breakdown in symbiotic fidelity is discussed in further detail in section 4.5 below, but was particularly associated with colonies inhabiting islands which have historically experienced severe and/or prolonged warming.

### 4.2. Expression adaptation and plasticity among coral and algal symbiont lineages

We used two approaches - variation partitioning and discriminant analysis of principal components (DAPC) - to assess the relative impact of the local environment, the primary genetic lineage, and the symbiotic partner on gene expression in both the *Pocillopora* host and the *Cladocopium* photosymbiont. Both analyses indicated a stronger influence of genetic lineage on photosymbiont gene expression than on the host (Figure 2B). Because we observed such high host-photosymbiont fidelity across the sampled colonies, it was difficult to identify which genetic factor (host or symbiotic partner lineage) was the primary driver of gene expression patterns. Given that the variation partitioning analysis indicated that for each partner (host and photosymbiont) their proper primary genetic lineage was the more influential genetic factor, we used those phylogenies, respectively, for informing our phylogenetic ANOVAs.

*Cladocopium C1* genes whose expression varied according to the photosymbiont lineage were enriched in PFam enrichments related to ion transport (57 genes) and papain family cysteine proteases (10 genes) which are known to show increased expression in plants in response to multiple environmental stressors (Liu et al. 2018a; Table S4). When grouped by genetic lineage of the host, *Cladocopium C1* gene expression profiles showed very little variation between colonies within groups suggesting a strong regulation of photosymbiont gene expression within a host lineage regardless of the environment (Figure 5B). In addition, photosymbiont expression profiles strongly mirrored the Pocillopora host phylogeny with Cladocopium from closely-related sister taxa in the host (e.g., SVD 3/5 and SVD 2/4) displaying more similar gene expression profiles than those inhabiting distantly related host lineages (Figure 5B). Again, this pattern would seem to suggest very strong, and potentially highly conserved, regulation of photosymbiont gene expression by the coral host.

Host genes whose variation in expression was principally attributed to the host genetic lineage were involved in biological processes related to regulation of proteolysis, including salvaging of amino acids, as well as genes involved in ‘cellular response to abiotic stimulus’, including ‘regulation of the innate immune response’, ‘defense response to virus’, and ‘response to ionizing radiation’ (GO:0071479, and GO:0010212; Table 2). This latter category was particularly enriched among genes overexpressed in host SVD1 present in the central Pacific.

By considering expression profile variation within its proper evolutionary context, we tested for evidence of expression-level adaptation among the five coral lineages identified in this study. We used the expression variance and evolution (EVE) model (Rohlfs and Nielsen 2015; Avila-Magaña et al. 2021), also referred to as Phylogenetic ANOVA, to identify genes with higher variance among than within lineages (divergent expression genes) and vice versa (diverse expression genes). The former set of genes display lineage-specific shifts in baseline expression suggestive of directional selection, potentially as a result of adaptive front-loading and/or altered gene copy number, and are therefore potential candidates for expression level adaptation. Conversely, the latter set of genes display higher expression variation within a lineage than among species suggesting either that regulation of these genes is highly conserved (i.e., low variation across lineages) and/or that these genes are highly responsive to local stimuli. These genes are therefore candidates for expression level conservation and/or plasticity.

In both partners we observed more divergent than diverse genes thus indicating that the majority of responsive genes displayed higher among-lineage gene expression variation than within-lineage differences. This further implies that although host and photosymbiont expression profiles differ significantly between lineages, within a lineage, profiles are relatively fixed. The influence of the genetic lineage is particularly strong for the Cladocopium photosymbiont in which very few genes appear linked to changes in the external environment (Figure 2B). We interpret this pattern as a signal that, in the photosymbiont, prior selection (i.e., adaptive mechanisms) appear to outweigh plastic responses to the local environment and that a given lineage of Cladocopium displays a gene expression profile that is largely “fixed” by its genotype with limited plastic potential. This genetic influence is also strong in the *Pocillopora* host (Figure 4A). However, host expression profiles were also more responsive to the local environment (Figures 2A and 3A) indicating that plastic response likely play a larger role in host acclimatization to the environment than in the photosymbiont.

### 4.3. Coral host transcriptomic plasticity may buffer the photosymbiont from environmental perturbation

Despite our finding that genotypic/adaptive effects may, in general, outweigh plastic responses among Pocillopora corals, we did observe a significant response to the environment among lineages. This was especially true for the coral host in which both the variation partitioning and DAPC approaches indicated a strong impact of the environment on coral host gene expression profiles.

Within the *Pocillopora* host, genes whose variation in expression was principally driven by differences in the local environment were involved in biological processes related to lipid metabolism/modification, cellular response to endogenous stimuli (especially growth factors), and signal transduction (Table 2). We interpret this differential sensitivity to the local environment as a signature of host-driven regulation of the photosymbiont micro-environment. Overall, we observed a reduced effect of the island of sampling on *Cladocopium C1* gene expression within a colony as compared to that of the coral host. These results also reflect those described by Leggat *et al.* (2011) who observed a significantly more robust transcriptional response in *Acropora aspera* coral hosts than in their associated photosymbionts following a simulated bleaching event. When exposed to acute heat stress, few photosymbiont genes showed significant shifts in expression level as compared to strong up-regulation of several host genes including heat shock proteins.

We interpret this combination of high host transcriptional plasticity and relatively low photosymbiont transcriptional variation as an indication of host-driven buffering of its hosted *Cladocopium* symbionts from environmental perturbation. Additionally, the high similarity we observed in *Cladocopium* expression profiles when hosted by closely-related *Pocillopora* lineages (Figure 5B) may further indicate strong regulatory control of photosymbiont expression by the host coral. Under this model, the majority of transcriptional acclimatization to the local environment is therefore driven by gene expression changes within the coral host. These responses likely serve to maintain symbiosome homeostasis and thereby ensure continued maintenance of host-specific photosymbiont expression states and continued regulation of photosymbiont productivity across environments. This also suggests that the primary mechanism of environmental adaptation in the photosymbiont is through selection of *Cladocopium* genotypes (i.e., high symbiont fidelity), either through active selection by the host or through winnowing of unfit populations within a colony, rather than through plastic, acclimatory, responses.

### 4.4. Elevated SST drives convergent expression among hosts and may select for heat-resistant algal genotypes

Constrained correspondence analysis of expression variation between islands, indicated that the strongest environmental factor contributing to colony gene expression was historical SST (Figure 6, Table S11). This is unsurprising given the primacy of temperature as an abiotic driver in ectothermic organisms generally (Somero et al. 2017) and given its specific, disruptory, effects on regulatory homeostasis within scleractinian corals (Hughes et al. 2018; Wall et al. 2021). In addition, discriminant analysis of principal components of host gene expression indicated that one of the two top discriminant functions was tightly linked to historical SST (DF2). This axis clearly distinguished colonies from Rapa Nui and Ducie islands (the coldest islands) from those inhabiting the Central Pacific (Gambier, Moorea, Aitutaki, and Niue, intermediate SSTs) and those inhabiting the the Western and Eastern Pacific (Upolu, Guam, Isla de Las Perlas, Coiba, and Malpelo, highest SSTs). Colonies from different genetic lineages and from geographically distant locations nevertheless displayed similar expression profiles along this discriminant axis, particularly when present on islands with historically elevated and/or low SSTs. This suggests a convergent plastic response to thermal challenge across *Pocillopora* lineages. Interestingly these same colonies also displayed high fidelity for three photosymbiont lineages - *Cladocopium* L2 and L5 as well as *Durusdinium D2* (Figure 1B).

In terms of coral host transcriptomic plasticity, we observed an up-regulation of genes traditionally associated with the environmental stress response (ESR) in colonies present on historically warm reefs. For example, host lineages from warm islands showed elevated expression of a 70-kDa heat shock protein, Heat Shock Protein Family A (Hsp70) Member 12A- like. This class of chaperones is known to be involved in response to thermal stress in scleractinian corals (Franzellitti et al. 2018). Because SSTs were not elevated on these islands at the time of sampling and because these colonies had not experienced an extreme heating event in the previous three weeks before sampling, we interpret this elevated expression of ESR genes among these colonies to be indicative of gene frontloading, a potential signal of local thermal adaptation. In addition, we observed enrichment among environmentally-responsive genes in xenobiotic transporters including both glutathione and lipid peroxidase activities. These functions may represent up-regulation of membrane-bound GSTs (MAPEG), an evolutionarily distinct class of enzymes that detoxify xenobiotic compounds and ameliorate oxidative stress (Kammerscheit et al. 2019).

In addition to transcriptomic plasticity within the cnidarian host, corals have many additional pathways permitting acclimatization to thermal challenges. At the colony level, thermal response integrates across the different components of the coral holobiont (coral host, algal photosymbiont, and associated microbiome/virome) which may respond at different timescale and/or through different physiological mechanisms. For example, whereas coral genotype is fixed within a colony, photosymbiont community composition can shift over time thereby altering the prevalence of the dominant genotype of the algal partner. Such shifts in the photosymbiont community can occur over short time scales, in response to acute thermal challenges for example (Claar et al. 2020), or over longer term periods as a result of competitive exclusion between photosymbiont lineages within a colony (Howells et al. 2020). Both mechanisms underpinning symbiont shift can indicate acclimatization of the coral holobiont to its local (thermal) environment.

In our dataset, we observed three photosymbiont lineages which appear to have been preferentially selected for in *Pocillopora* colonies inhabiting islands with historically elevated SSTs. Whereas non-selective mechanisms could account for this pattern. Dispersal limitations can result in the dominance of particular photosymbionts on certain reefs, particularly if those reefs are geographically isolated. However, dispersal limitation alone seems insufficient as an explanation for the high photosymbiont specificity we observed in this study in part because we observe the same photosymbiont communities present on reefs ca. 14,000 km apart across the Pacific basin (e.g., Coiba and Guam). The high holobiont fidelity we observed under elevated SSTs may therefore truly represent a signature of differential selection in response to local thermal conditions (i.e., local thermal adaptation).

Although we did not directly assess photosymbiont or colony fitness across environments in this study, we did examine holobiont gene expression profiles which provide some context regarding the relative physiological states of colonies within a given environment. For example, in the symbiotic partner DAPC model, one of the genes which significantly discriminated between *Pocillopora* colonies containing different *Cladocopium C1* photosymbionts was a DNAJ chaperone protein. This gene was also among the candidate adaptive genes whose expression was highly divergent across *Pocillopora* lineages. We observed reduced expression of this protein in colonies from the coldest island (Rapa Nui). However, we also observed elevated expression of this protein in colonies containing *Cladocopium* L2 symbionts in the Central Pacific (all SVD 1 corals on Ducie, Gambier, and Moorea, Figure 1B and C). Interestingly this gene was only upregulated on these islands in SVD1/L2 holobiont colonies; SVD4/L4, SVD2/L5, and SVD5/L3 holobiont colonies from the same islands did not exhibit similar upregulation of this chaperone protein. This gene was also not strongly upregulated among SVD5/L3 colonies from islands in the Western Pacific that have historically experienced much higher SSTs (and which were warmer at the time of sampling) than those of the Central Pacific. This indicates that while this DNAJ chaperone protein may play an important role in thermal acclimatization in the SVD1/L2 Pocillopora holobiont, it is not a significant contributor to thermal acclimatization in the SVD5/L3 holobiont.

In addition, although they represent only a small fraction of the top variant genes, *Pocillopora* genes with expression dependent on the symbiont lineage were involved in several important biological processes including those potentially related to regulation of calcification, symbiont photosynthesis, and to the scavenging of reactive oxygen species (Table 2). For example, genes with expression variation linked to symbiont lineage were highly enriched in functions related to transport of ions across cellular membranes including ATPase-coupled transmembrane transporters which are known to play a role in host calcification (Barott et al. 2015b, 2020; Zoccola et al.) and as carbon-concentration mechanisms in marine photosymbioses thereby regulating *Symbiodiniaceae* photosynthesis (2015a, b; Tresguerres et al. 2017; Armstrong et al. 2018).

Finally, as discussed in section 4.2 previously, the low expression variation we observed between Cladocopium lineages inhabiting warm reefs may itself reflect convergent evolution in response to environmental selection. Photosymbiont expression profiles may be less affected by the local environment simply because only certain symbiont genotypes are found in warm environments. That is to say, algal expression profiles are similar because of shared ancestry/symbiont fidelity due to historical selection for heat-tolerant symbiotypes. When taken together, the strong correlation between historical SST and photosymbiont community composition, the presence of similar algal genotypes on extremely distant reefs, as well as the reduced expression of host chaperone proteins relative to colonies containing other symbiont communities all strongly suggest the presence of a selective signature in these photosymbiont lineages, thereby implying environmental specialization (i.e., local adaptation) for increased heat tolerance.

### 4.5. Idiosyncratic host expression profiles associated with symbiont shuffling under thermal challenge

For example, both *Cladocopium-* and *Durusdinium-*dominant *Pocillopora* colonies were found in the Eastern and Western Pacific and this appearance coincides with historically elevated SST at those reefs (see section 3.5 below). One interpretation of these observations is that they provide evidence for thermally-driven symbiont switching/shuffling in these colonies (e.g., shuffling of dominant symbionts or recolonization after bleaching). However, because *Pocillopora meandrina* symbionts are generally acquired via vertical transmission, with oocytes being “seeded” with the maternal symbiont community (Marlow and Martindale 2007), it is also possible that these C- and D-containing colonies represent true, evolutionarily stable, holobiotypes which persist across generations on reefs which experience frequent and/or severe thermal stress.

The known switch between the thermo-sensitive *Cladocopium* to the thermo-tolerant *Durusdinium* (LaJeunesse et al. 2010), observed here mainly in the eastern Pacific, suggests that *Pocillopora* colonies may have recently experienced acute heat stress just prior to sampling.

In the Eastern Pacific, we observed both *Cladocopium*- and *Durusdinium*-dominant colonies of Pocillopora SVD 4. These colonies were present on reefs which have historically experienced frequent, extreme thermal stress providing evidence for ‘symbiont switching’ as a thermal acclimation strategy in this host. However, symbiont switching had very little effect on host gene expression *in situ* and host expression appears driven more by the environment (both at the site and island scale) than by symbiont lineage. Our data for *Pocillopora* show that switching between *Cladocopium* and *Durusdinium* algal communities had only a weak, and inconsistent, impact on host expression *in situ*. This stands in contrast to previous studies which have reported a significant role of the photosymbiont in modulating coral host gene expression (DeSalvo et al. 2010). For example, a recent study in *Montastraea cavernosa* reported that colonies containing *Durusdinium* symbionts displayed elevated expression of ESR genes even under ambient conditions relative to colonies containing *Cladocopium* symbionts (Cunning and Baker 2020). These results suggest that symbiotype significantly alters host expression even prior to environmental perturbation.

On average, we found 129 genes differentially expressed between C- and D-containing Pocillopora colonies in the Eastern Pacific (Table S2). In total, only 4% of gene expression variation between Eastern Pacific coral colonies was attributed to differences in symbiotype and this was not significant (PERMANOVA, p > 0.05). In contrast, in *M. cavernosa* colonies, up to 14% of expression variation between individuals could be attributed to the switch between dominant symbiotypes (Cunning and Baker 2020). In our dataset, the environment played a much more significant role in determining *Pocillopora* gene expression with 24% of expression variation being explained by sampling island and, unlike symbiotype, reef environment contributed significantly to overall host gene expression in these colonies (PERMANOVA, p = 0.001; Figure S5). Additionally we observed very little to no overlap in differentially expressed genes between C- and D-containing Pocillopora colonies among the three Eastern Pacific islands (Figure S6). This implies that there was no common influence of *Durusdinium* symbionts on host expression across reefs. Finally, we observed high gene expression variation among *Durusdinium*-containing *Pocillopora* colonies from within the same reef (Figures S5 and S6) suggesting that, even at small spatial scales, other factors were more important than symbiotype in determining host gene expression.

One explanation for the greater similarity in gene expression between C- and D-containing colonies in *Pocillopora* SVD 4 relative to *M. cavernosa* may be related to the influence of both the local thermal environment and variation in host genotype on host expression in our dataset. We observed D-dominant Pocillopora colonies only on reefs which have historically experienced frequent, severe heating events (Isla de las Perlas, Coiba, Malpelo, and Guam). We therefore suggest that both C- and D-containing colonies displayed elevated expression of ESR genes on these reefs explaining why we don’t observe the specific, ‘thermal priming’ influence of *Durusdinium* on the host. In contrast, the *M. cavernosa* colonies sampled by Cunning and Baker (2020) were reared under controlled conditions and represented ramets of the same genet. These controlled factors in the *M. cavernosa* colonies may have allowed for maximization of the ‘symbiotype signal’ relative to the environmental and genetic influences on host gene expression. Thus, it’s possible that the similarity in host expression we observed in *Pocillopora* between C- and D-containing colonies is a result of physiological canalization (e.g., a universal ESR) in the host which partially obscured the putative front loading of these genes in colonies containing *Durusdinium* as observed in *M. cavernosa* under controlled conditions.

Despite the weak influence of the dominant symbiotype on overall *Pocillopora* host expression in the Eastern Pacific, we did observe a small set of genes that were differentially expressed between C- and D-containing hosts. These included genes associated with ‘chaperone-mediated protein folding’ (GO:0061077), ‘protein ubiquitination’ (GO:0016567), and ‘regulation of apoptosis’ (GO:0042981) which were among the most significantly enriched biological process terms associated with genes down-regulated in D-containing colonies relative to C-containing hosts in some reefs (Table S2). These results provide indirect support for the Adaptive Bleaching Hypothesis in that coral colonies containing the more thermally tolerant *Durusdinium* symbiont show reduced expression of ESR as compared to colonies containing *Cladocopium* symbionts inhabiting the same reef.

### 4.6. Three-tiered strategy of thermal acclimatization in Pocillopora holobionts

When examined together, our metagenomic and metatranscriptomic datasets seem to imply the existence of three, potentially sequential, strategies for thermal acclimatization in *Pocillopora* holobionts: 1) selection of and high fidelity for heat tolerant Cladocopium lineages in warm environments, 2) high host transcriptomic plasticity which buffers photosymbiont against thermal challenge, and 3) symbiont shuffling and replacement of *Cladocopium* with *Durusdinium*. However, the effectiveness of these strategies or their relative thresholds of initiation may differ between *Pocillopora* lineages. For example, we observed a general trend towards symbiont fidelity among Pocillopora lineages (Figure 1B) and this fidelity appears linked to local sea surface temperatures. Similar high host-photosymbiont fidelity has also been reported in corals inhabiting some of the hottest reefs in the world in the Persian Gulf (Howells et al. 2020). This tendency for high symbiotic fidelity may therefore represent the first step in a strategy of thermal acclimatization in the coral metaorganism whereby heat tolerant photosymbiont lineages are preferentially selected/maintained under elevated SST.

Despite the trend towards high symbiont fidelity, we did observe instances of symbiont flexibility in SVD 4 and SVD 5 hosts. In both cases, this flexibility was characterized by the transition of photosymbiont communities towards two putatively heat tolerant lineages (*Cladocopium C1* L3 and L5) suggesting that some *Pocillopora* lineages can initiate plastic shifts in symbiont community composition as a potential strategy for coping with environmental warming. However, the physiological effects of this symbiont flexibility may not be equivalent among lineages. For example, when present on the coldest island (Rapa Nui) host SVD 5 corals showed strong fidelity with *Cladocopium* L1. However, when present on warmer islands in the central Pacific, SVD 5 corals were always found in symbiosis with *Cladocopium* L3, a putative heat tolerant lineage. Despite this shift in symbiont community, however, host expression of a DNAJ chaperone protein was extremely elevated in L3-containing SVD 5 colonies relative to other lineages in the same environments. Conversely, when present in the central Pacific, host SVD 4 colonies showed strong fidelity with *Cladocopium* L4 and exhibited relatively high DNAJ expression. DNAJ was down-regulated, however, when this same lineage was present on some of the warmest reefs sampled in this study (eastern Pacific) where the SVD 4 corals were found in symbiosis with another putatively thermo-tolerant photosymbiont, *Cladocopium* L5. Although the high DNAJ expression observed in SVD 5 corals in the central Pacific could represent a positive acclimatization response (i.e., gene frontloading) accompanying the symbiont flexibility, it could also be indicative of the onset of thermal stress in these colonies. In this case, the high DNAJ expression we observed in SVD 5 colonies relative to SVD 4 colonies present on warmer reefs may signal that the SVD 5 holobionts are already at or near their upper heat tolerance limits and may therefore only be able to rely on symbiont shuffling to resist future warming. Evidence from other coral species, however, suggests that symbiont shuffling may only be effective as a “last ditch” strategy to weather acute extreme heating events and that this strategy, while increasing survival in the short-term, may negatively impact resilience in colonies under future warming events (Claar et al. 2020).

## 5. CONCLUSION

Overall, we observed high symbiotic fidelity among *Pocillopora* lineages across environments with evidence of potential selection for heat-resistance photosymbiont lineages on islands with historically elevated sea surface temperatures. We reveal that host gene expression profiles are strongly segregated by genetic lineage and environment, and were significantly correlated with several historical sea surface temperature (SST) traits. In contrast, *Cladocopium C1* expression profiles were primarily driven by algal genotype and displayed low phenotypic plasticity across environmental gradients. Overall, our data suggest the existence of a three-tiered strategy underpinning thermal acclimatization in *Pocillopora* holobionts with strong selection for specific photosymbiont lineages in certain environments (i.e., host-photosymbiont fidelity), high host transcriptomic plasticity acting as an environmental buffer, and a ‘last ditch’ strategy of photosymbiont shuffling playing progressive roles in response to elevated SSTs. Our study has important implications for continued coral reef conservation and prediction of coral responses to future ocean warming. Our examination of the mechanisms underpinning acclimatization among lineages across an environmental gradient and our identification of potential candidate loci under thermal selection provide useful baseline metrics for informing future manipulative experiments focused on physiological mechanisms underlying these observations. Investigating the extent to which different thermal acclimatization strategies, including both transcriptional and compositional changes, are used among lineages will allow us to better manage coral reefs under future ocean warming.

---

**Supplemental Figure 1.**
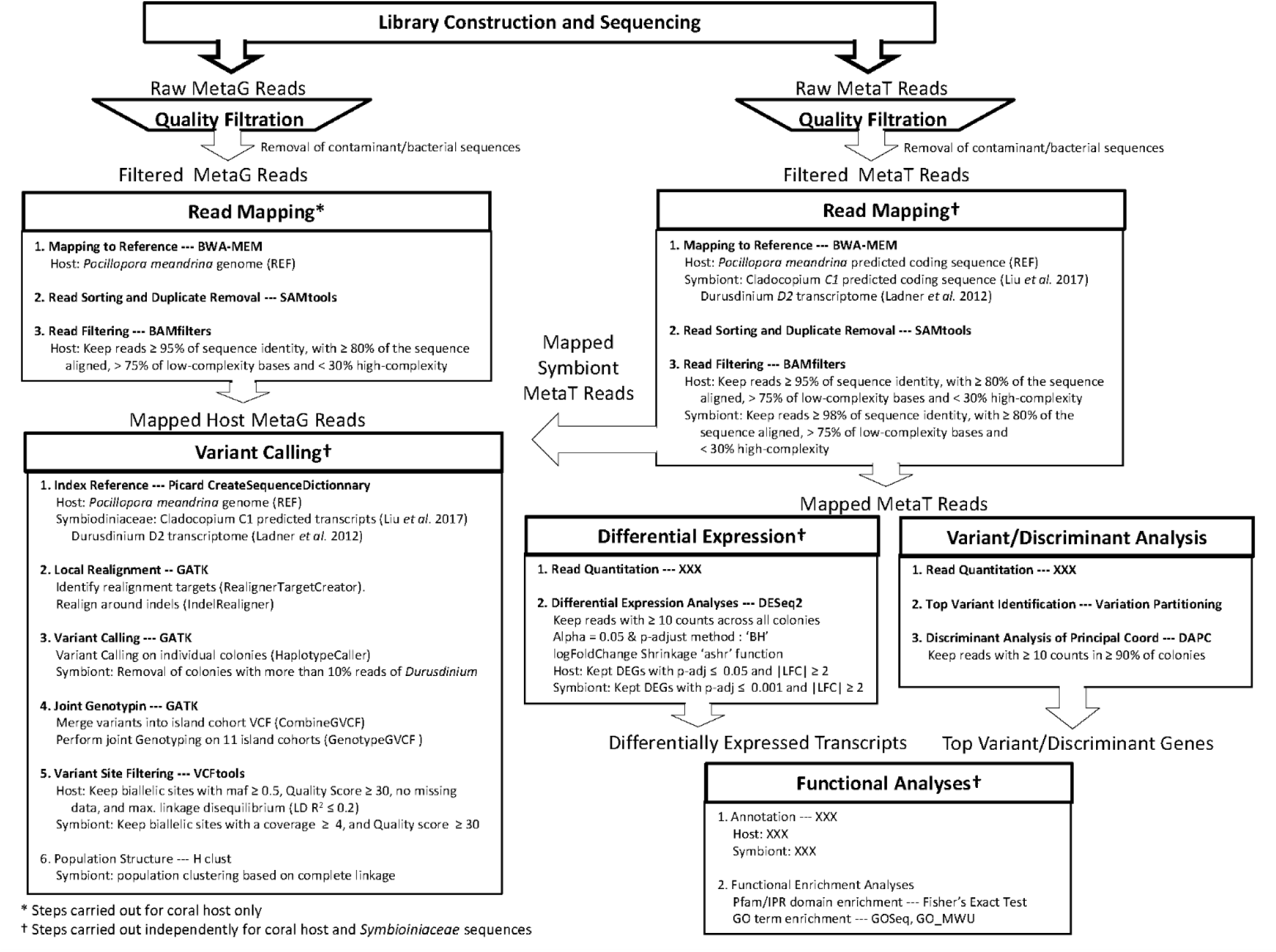
Bioinformatic workflow. Summary of the bioinformatic workflow used to identify single nucleotide polymorphisms (SNPs) in the host and symbiont as well as to perform functional analyses.

**Supplemental Figure 2.**
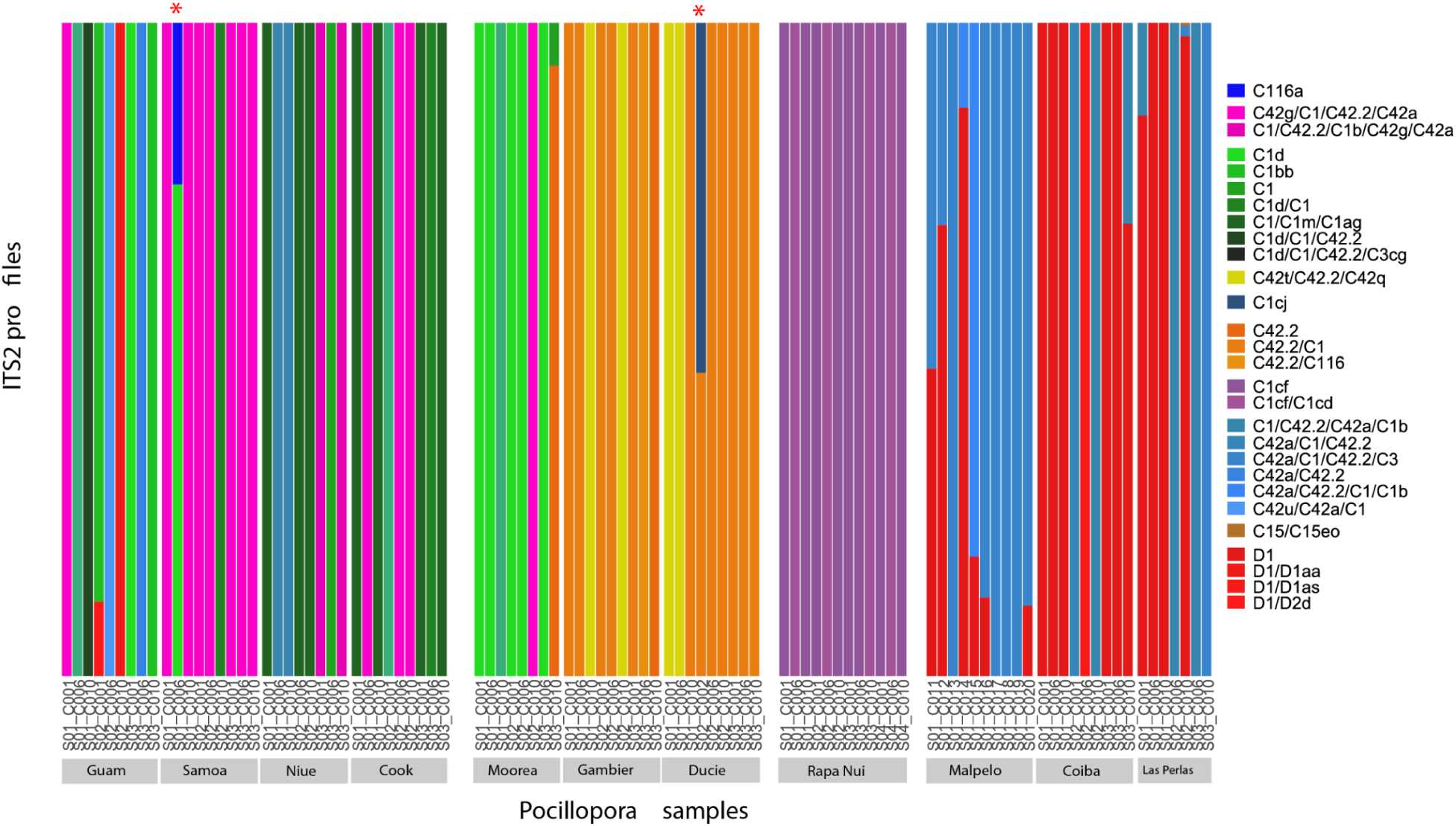
Relative abundance of Symbiodiniaceae ITS2 profiles in Pocillopora colonies. The abundance of each ITS2 profile was obtained from ITS2 reads analyzed with the SymPortal method (Hume *et al.* 2019). Each color corresponds to a different profile named by its most abundant ITS2 sequence. *Pocillopora* colonies indicated by an asterisk (top) contain 2 different *Cladocopium* ITS2 profiles and were removed from the population structure analysis.

**Supplemental Figure 3.**
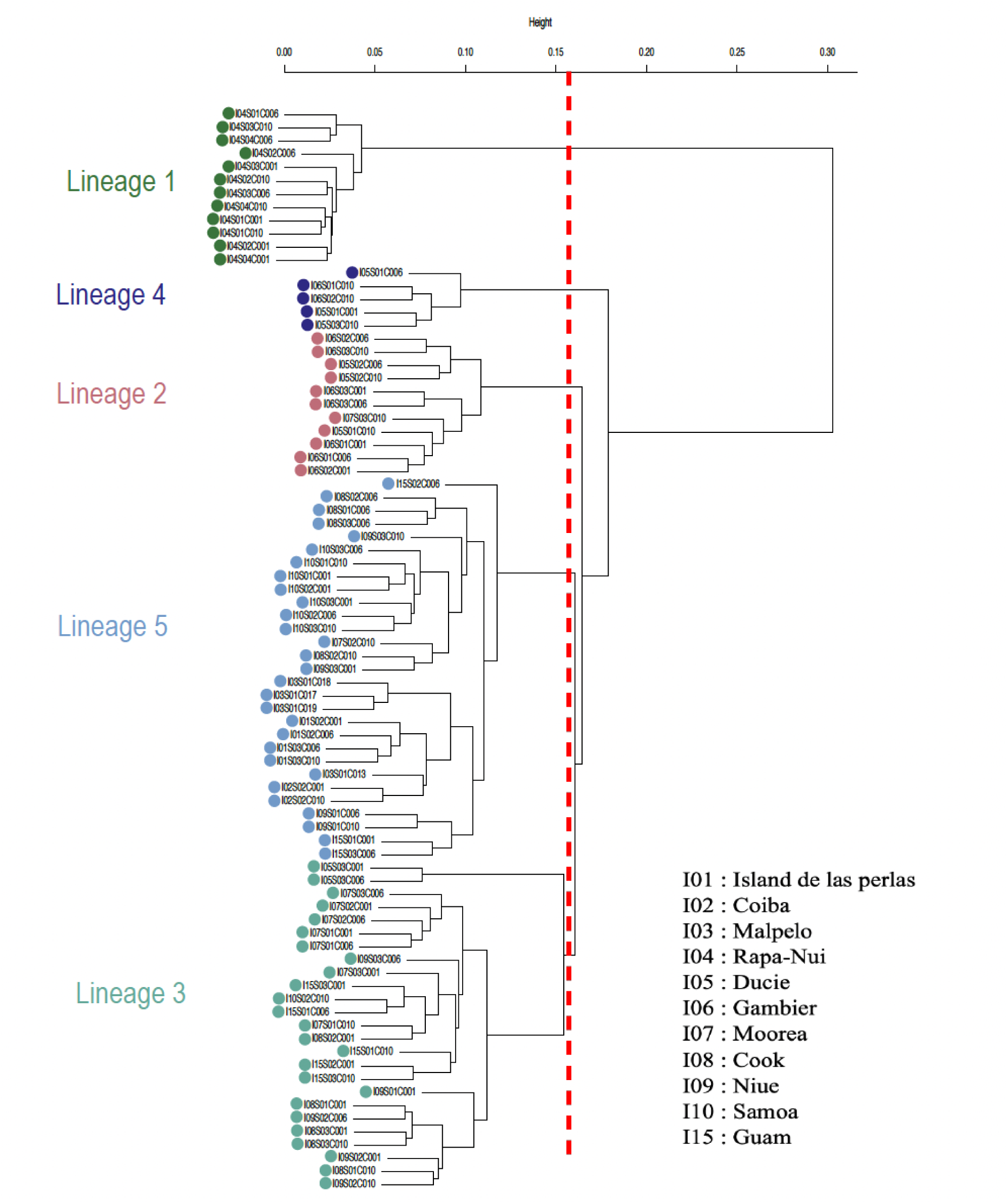
Hierarchical clustering of Cladocopium C1 populations in metatranscriptomic samples of Pocillopora spp. Dendrogram representation of the hierarchical clustering of 3.712 SNPs detected by the mapping of metatranscriptomic reads on 1,354 genes of the *Cladocopium* C1 reference genome (Liu et al. 2018b). 82 *Cladocopium* samples were clustered into 5 genetic lineages based on a tree height cutoff threshold (red line).

**Supplemental Figure 4:**
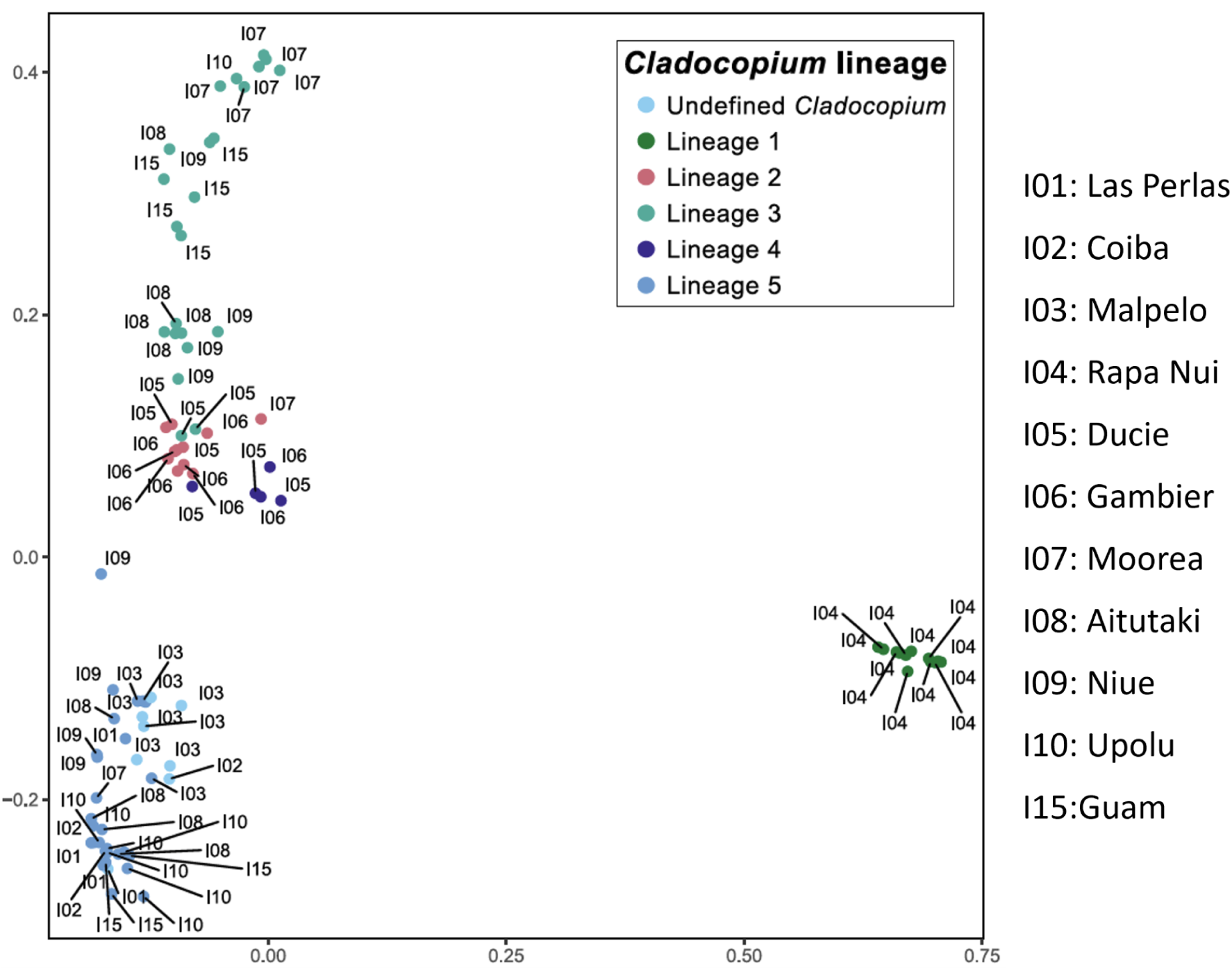
Cladocopium ITS2 profiles distances compared to SNP-based clustering. ITS2 profiles of *Pocillopora*-containing *Cladocopium* colonies were analyzed using the SymPortal method (Hume *et al.* 2019). Unifrac distances between each pair of samples were calculated and are represented in a principal coordinate analysis (PCA). Each dot represents a *Pocillopora* colony coloured according to the *Cladocopium C1* lineage determined by the SNP frequencies on the genomic coding sequence.

**Supplemental Figure 5.**
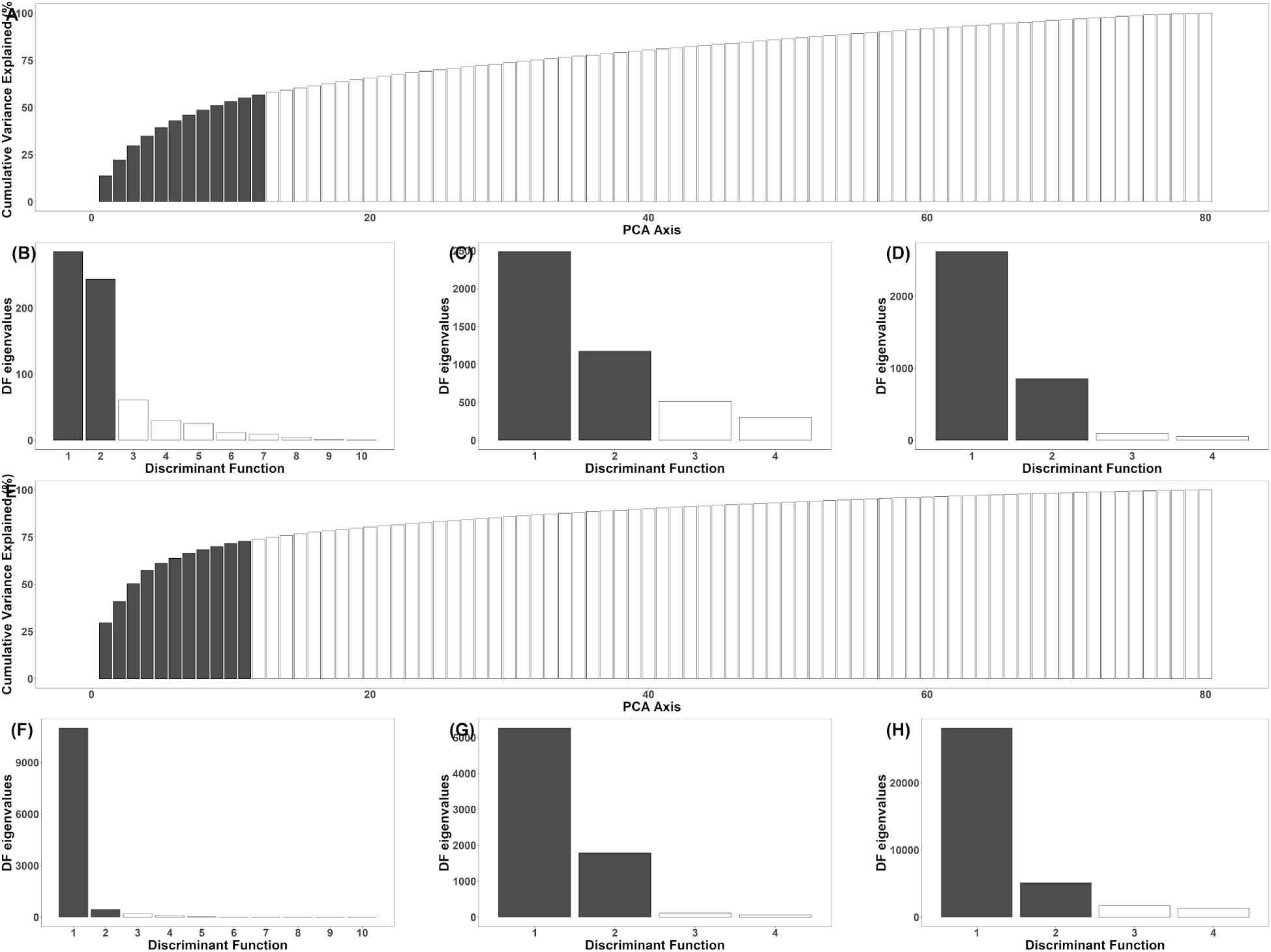
DAPC Eigenvalues. Cumulative percentage of variance explained by the principal coordinate axes (PCA) of the DAPC model for the host (A) and photosymbiont (E). Shaded bars indicate the PCAs retained in each model respectively. Discriminant function eigenvalues for the DAPC models grouped by the environment (B,F), the primary lineage (C,G), and the lineage of the symbiotic partner (D,H) for the host and symbiont, respectively.

**Supplemental Figure 6.**
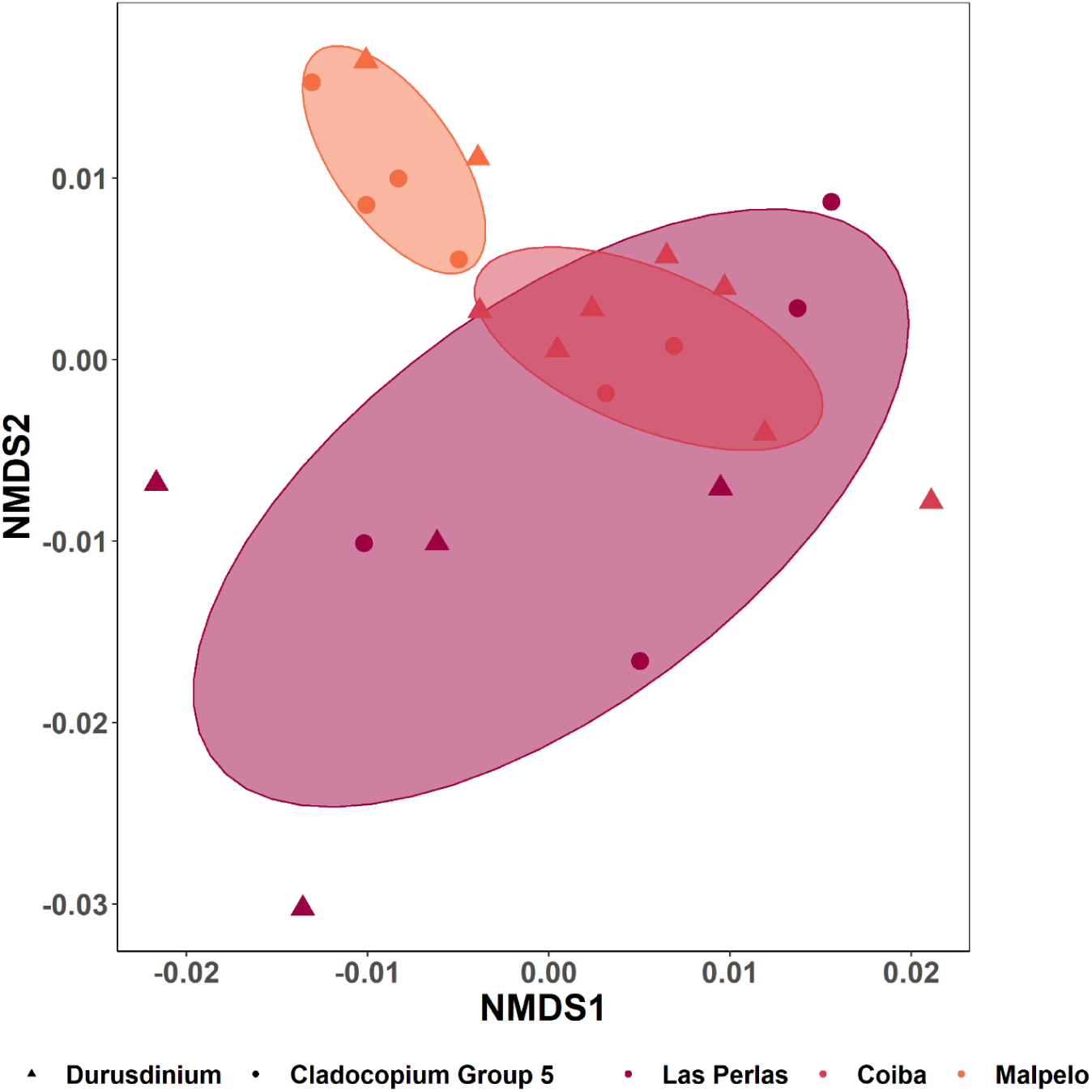
MDS plot of global host gene expression profiles of Cladocopium- and Durusdinium-containing Pocillopora SVD 4 colonies in the Eastern Pacific.

**Supplemental Figure 7.**
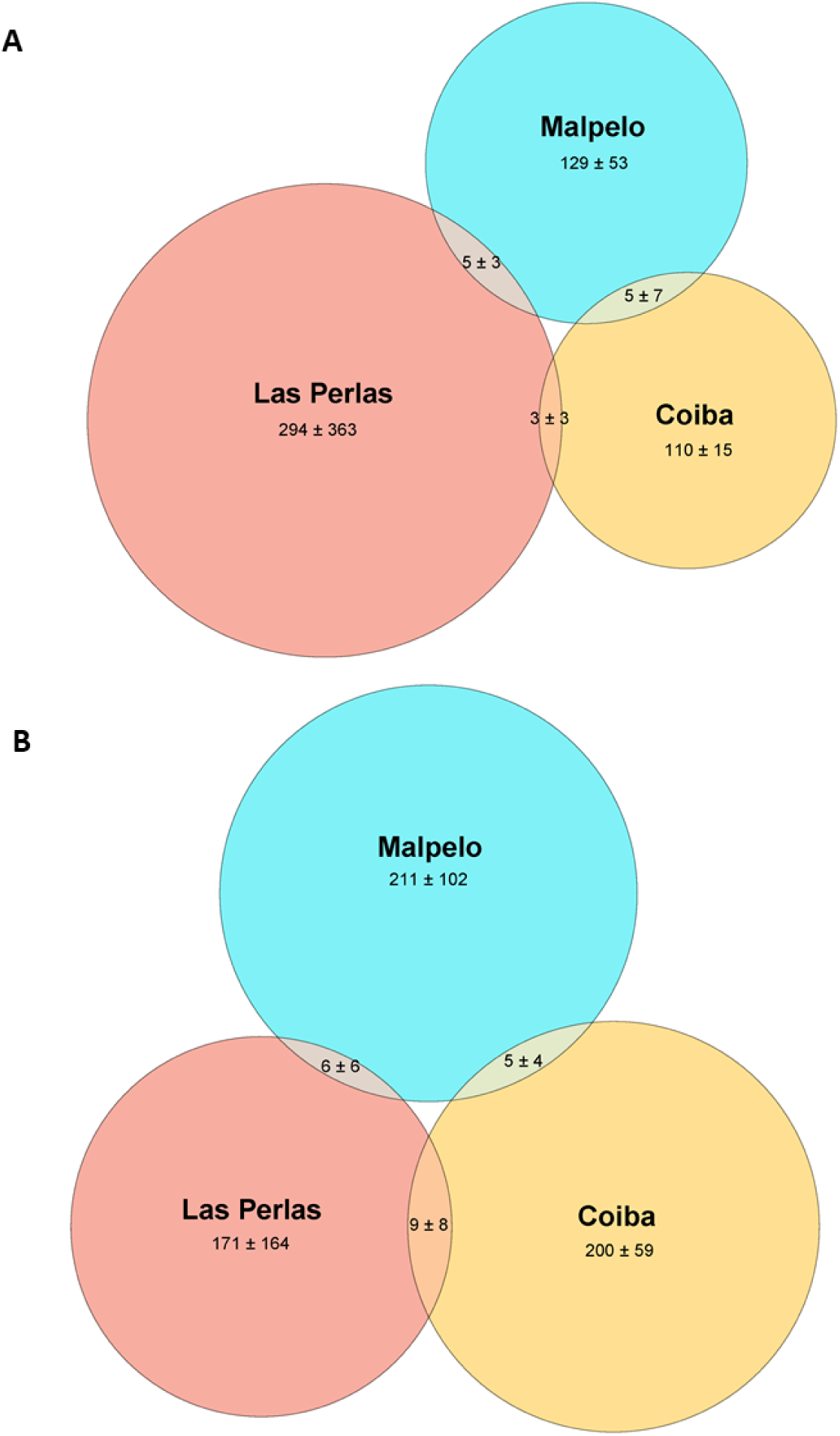
Summary of host genes differentially expressed between Cladocopium- and Durusdinium-containing Pocillopora SVD 4 colonies in the Eastern Pacific. Mean number (± sd) of *Pocillopora* host genes (A) up- and (B) down-regulated in D-containing colonies relative to those with *Cladocopium* after controlling for discrepant colony numbers on three islands in the Eastern Pacific (Isla de las Perlas, Coiba, and Malpelo).

**Supplemental Figure 8.**
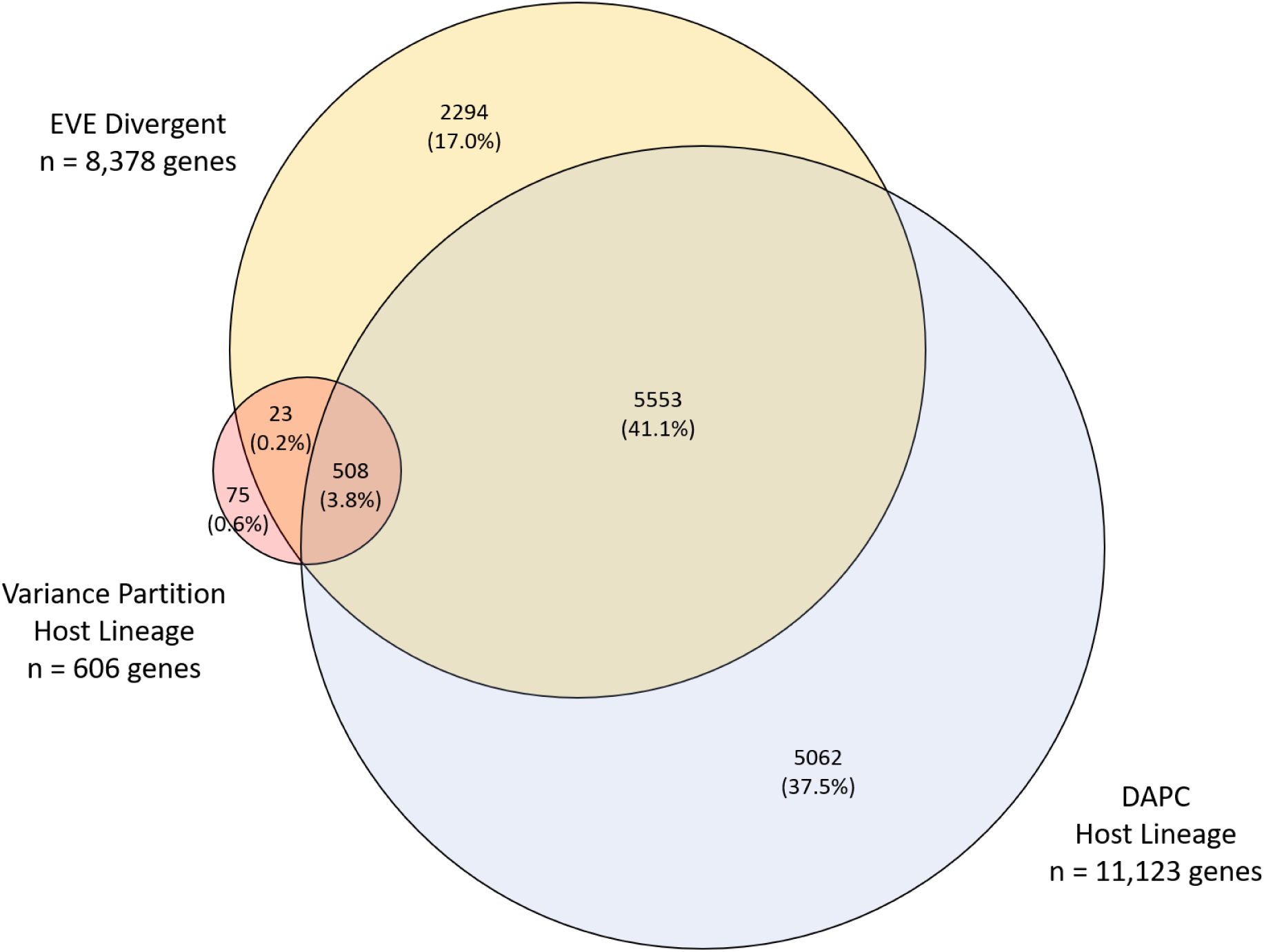
Overlap among top variant (red), divergent (yellow), and discriminant (blue) genes in the Pocillopora coral host.

**Supplemental Figure 9.**
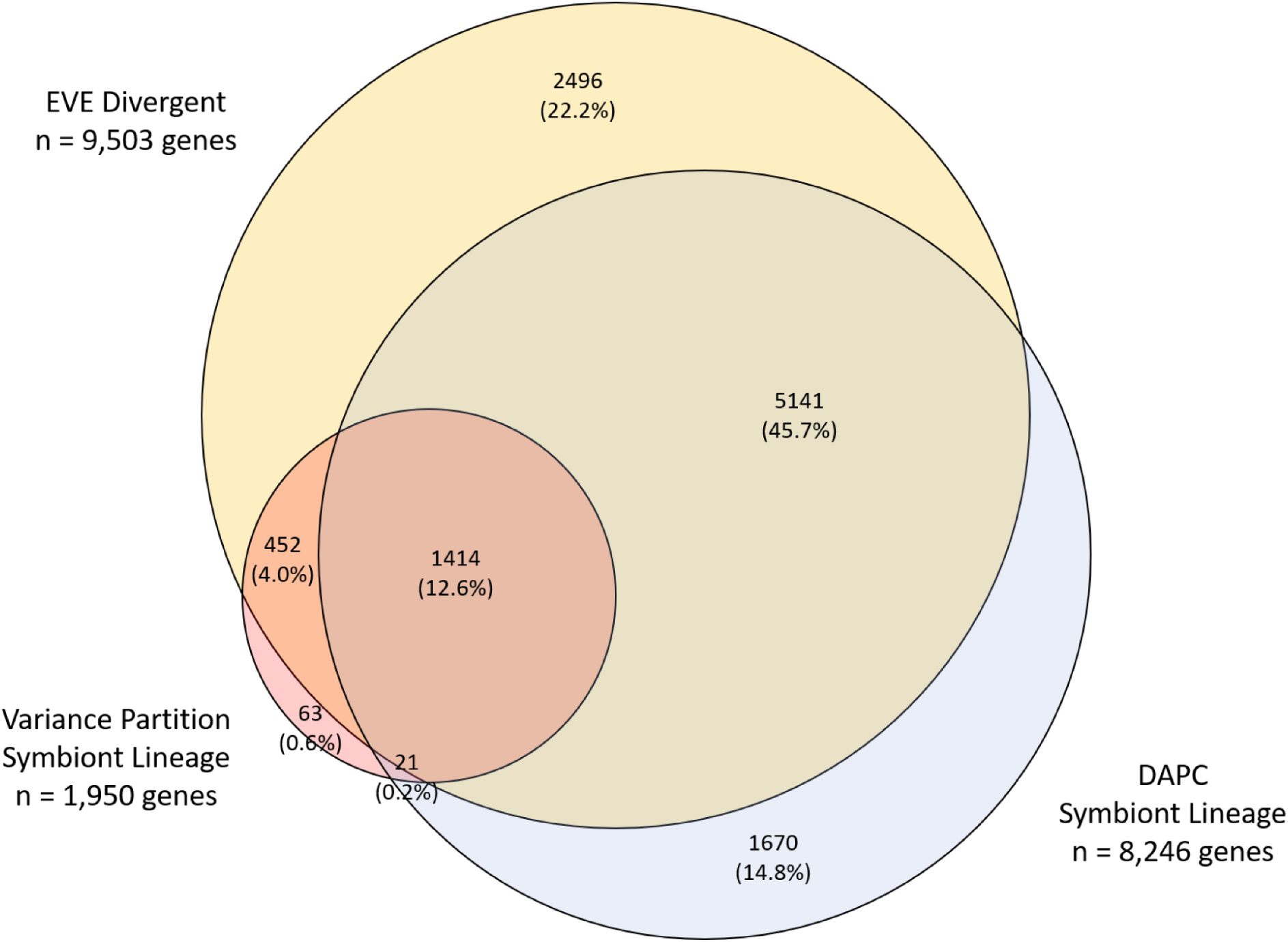
Overlap among top variant (red), divergent (yellow), and discriminant (blue) genes in the Cladocopium photosymbiont.

## TABLES

*Supplemental Table 1.*

*Summary of biological process functional enrichments among environment-associated top variant and discriminant genes in Pocillopora.*

*See supplemental files.*

*Supplemental Table 2.*

*Summary of functional enrichments among environment-associated top variant and discriminant genes in Cladocopium.*

*See supplemental files.*

*Supplemental Table 3.*

*Summary of biological process functional enrichments among primary genetic lineage-associated top variant and discriminant genes in Pocillopora.*

*See supplemental files.*

*Supplemental Table 4.*

*Summary of biological process functional enrichments among primary genetic lineage-associated top variant and discriminant genes in Cladocopium.*

*See supplemental files.*

*Supplemental Table 5.*

*Summary of biological process functional enrichments among symbiotic partner-associated top variant and discriminant genes in Pocillopora.*

*See supplemental files.*

*Supplemental Table 6.*

*Summary of biological process functional enrichments among symbiotic partner-associated top variant and discriminant genes in Cladocopium.*

*See supplemental files.*

**Supplemental Table 7.** Number of differentially expressed genes (p <= 0.05 or 0.001 in the host and symbiont, respectively; |LFC| >= 2) recovered in all intra-lineage, inter-island pairwise comparisons in Pocillopora spp.

| Holobiont | Contrast | Organism | DEGs (N) |
| --- | --- | --- | --- |
| SVD 1 - SymGG 2 | Ducie vs Gambier | Host | 409 |
|  |  | Symbiont | 377 |
| SVD 2 - SymGG 5 | Aitutaki vs Niue | Host | 95 |
|  |  | Symbiont | 22 |
|  | Aitutaki vs Upolu | Host | 195 |
|  |  | Symbiont | 1545 |
|  | Aitutaki vs Guam | Host | 266 |
|  |  | Symbiont | 329 |
|  | Niue vs Upolu | Host | 94 |
|  |  | Symbiont | 1357 |
|  | Niue vs Guam | Host | 322 |
|  |  | Symbiont | 264 |
|  | Upolu vs Guam | Host | 402 |
|  |  | Symbiont | 216 |
| SVD 3 - SymGG 3 | Aitutaki vs Niue | Host | 505 |
|  |  | Symbiont | 28 |
|  | Aitutaki vs Guam | Host | 384 |
|  |  | Symbiont | 686 |
|  | Niue vs Guam | Host | 291 |
|  |  | Symbiont | 481 |
| SVD 4 - SymGG 4 | Ducie vs Gambier | Host | 572 |
|  |  | Symbiont | 201 |
| SVD 4 - SymGG 5 | Las Perlas vs Coiba | Host | 322 |
|  |  | Symbiont | 71 |
|  | Las Perlas vs Malpelo | Host | 629 |
|  |  | Symbiont | 1556 |
|  | Coiba vs Malpelo | Host | 268 |
|  |  | Symbiont | 302 |
| SVD 5 - SymGG 3 | Ducie vs Moorea | Host | 556 |
|  |  | Symbiont | 1541 |

Total = 5,310 host DEGs, 4,597 symbiont DEGs.

**Supplemental Table 8.** Number and functional enrichment of differentially expressed host genes (p <= 0.05 and |LFC| >= 2) recovered in all intra-island comparisons between Cladocopium- and Durusdinium- containing Pocillopora SVD 4 colonies in the Eastern Pacific.

| Comparison | Colonies (C/D; N) | Category | DEGs (N) | Enriched BP Terms (GO) |
| --- | --- | --- | --- | --- |
| *Pairwise Intra-island* |  |  |  |  |
| Isla de las Perlas | 4 / 4 | Up-regulated in D | 69 | Bioluminescence (GO:0008218)<br>Xenobiotic transport (GO:0042908)<br>Gluconeogenesis (GO:0006094)<br>Mitochondrial protein catabolic process (GO:0035694)<br>Base-excision repair (GO:0006284)<br>Transmembrane transport (GO:0055085) |
|  |  | Down-regulated in D | 53 | Calcium ion transmembrane transport (GO:0070588)<br>Phospholipid biosynthetic process (GO:0008654) |
| Coiba | 2 / 7 | Up-regulated in D | 51 | Protein phosphorylation (GO:0006468)<br>Proteolysis (GO:0006508) |
|  |  | Down-regulated in D | 15 | Chaperone-mediated protein folding (GO:0061077)<br>(Obsolete) chaperone-mediated protein transport (GO:0072321)<br>Synaptic vesicle transport (GO:0048489)<br>Nuclear envelope organization (GO:0006998)<br>Protein phosphorylation (GO:0006468)<br>Cell adhesion (GO:0007155)<br>Neuron projection development (GO:0031175)<br>Intermediate filament cytoskeleton organization (GO:0045104) |
| Malpelo | 4 / 2 | Up-regulated in D | 120 | Bioluminescence (GO:0008218)<br>Regulation of signaling receptor activity (GO:0010469)<br>Vacuolar transport (GO:0007034)<br>Xenobiotic transport (GO:0042908) |
|  |  | Down-regulated in D | 276 | Bioluminescence (GO:0008218)<br>Regulation of apoptotic process (GO:0042981)<br>Protein phosphorylation (GO:0006468)<br>S-adenosylmethionine biosynthetic process (GO:0006556)<br>Regulation of pH (GO:0006885)<br>Notch signaling pathway (GO:0007219)<br>MyD88-dependent toll-like receptor signaling pathway (GO:0002755)<br>Proteolysis (GO:0006508)<br>Signal transduction (GO:0007165)<br>Protein ubiquitination (GO:0016567)<br>Netrin-activated signaling pathway (GO:0038007) |
| *Combined Across Environments* |  |  |  |  |
| All Together | 10 / 13 | Up-regulated in D | 109 | Anion transport (GO:0006820)<br>Response to pheromone (GO:0019236)<br>SCF-dependent proteasomal ubiquitin-dependent protein catabolic process (GO:0031146)<br>Positive regulation of canonical Wnt signaling pathway (GO:0090263)<br>Negative regulation of apoptotic process (GO:0043066)<br>GPI anchor biosynthetic process (GO:0006506)<br>Immune response (GO:0006955) |
|  |  | Down-regulated in D | 20 | MyD88-dependent toll-like receptor signaling pathway (GO:0002755)<br>Positive regulation of I-kappaB kinase/NF-kappaB signaling (GO:0043123) |

**Supplemental Table 9.** Discriminant analysis of principle component (DAPC) model information. Information regarding the number of principal components (PC) and discriminant functions (DF) retained, the proportion of gene expression variance explained (Var), and overall mean correct reassignment proportions (Assign) of the six discriminant analysis of principal component (DAPC) models.

| **Model** | **Species** | **PC** | **DF** | **Var** | **Assign** |
| --- | --- | --- | --- | --- | --- |
| Environment | *Pocillopora* | 12 | 10 | 0.567 | 0.963 |
|  | *Cladocopium C1* | 11 | 10 | 0.728 | 0.914 |
| Primary Lineage | *Pocillopora* | 12 | 4 | 0.567 | 1.000 |
|  | *Cladocopium C1* | 11 | 4 | 0.728 | 1.000 |
| Symbiotic Partner | *Pocillopora* | 12 | 4 | 0.567 | 0.988 |
|  | *Cladocopium C1* | 11 | 4 | 0.728 | 0.988 |

**Supplemental Table 10.** Permutational multivariate analysis of variance (PERMANOVA) model information. Table containing the results of permutational multivariate analysis of variance (PERMANOVA) for Pocillopora and Cladocopium C1 gene expression under the three a priori single factor grouping scenarios (environment, primary lineage, and symbiotic partner) as well as for the interactive model (environment x primary genetic lineage). Values include the model degrees of freedom (DF), the sum (Sum Sq) and mean (Mean Sq) of squares, the model F value for the grouping factor, and the correlation coefficient (R^2^).

| Model | Species | Factor | DF | Sum Sq | Mean Sq | Model F | $R^2$ |
| --- | --- | --- | --- | --- | --- | --- | --- |
| Environment | *Pocillopora* | Island | 10 | 571332 | 57133 | 4.1688 | 0.373 *** |
|  |  | Residuals | 70 | 959335 | 13705 |  | 0.627 |
|  |  | Total | 80 | 1530667 |  |  | 1.000 |
|  | *Cladocopium C1* | Island | 10 | 240829 | 24082.9 | 9.2265 | 0.569 *** |
|  |  | Residuals | 70 | 182713 | 2610.2 |  | 0.431 |
|  |  | Total | 80 | 423542 |  |  | 1.000 |
| Primary Lineage | *Pocillopora* | SVD Clade | 4 | 384787 | 96197 | 6.3802 | 0.251 *** |
|  |  | Residuals | 76 | 1145880 | 15077 |  | 0.749 |
|  |  | Total | 80 | 1530667 |  |  | 1.000 |
|  | *Cladocopium C1* | Lineage | 4 | 215284 | 53821 | 19.641 | 0.508 *** |
|  |  | Residuals | 76 | 208258 | 2740 |  | 0.492 |
|  |  | Total | 80 | 423542 |  |  | 1.000 |
| Symbiotic Partner | *Pocillopora* | Lineage | 4 | 378529 | 94632 | 6.2423 | 0.247 *** |
|  |  | Residuals | 76 | 1152138 | 15160 |  | 0.753 |
|  |  | Total | 80 | 1530667 |  |  | 1.000 |
|  | *Cladocopium C1* | SVD Clade | 4 | 151960 | 37990 | 10.631 | 0.359 *** |
|  |  | Residuals | 76 | 271582 | 3573 |  | 0.641 |
|  |  | Total | 80 | 423542 |  |  | 1.000 |
| Interactive | *Pocillopora* | Island | 10 | 571332 | 57133 | 4.8836 | 0.373 *** |
|  |  | SVD Clade | 4 | 166238 | 41559 | 3.5524 | 0.109 *** |
|  |  | Island:SVD Clade | 5 | 79459 | 15892 | 1.3584 | 0.052 ** |
|  |  | Residuals | 61 | 713638 | 11699 |  | 0.466 |
|  |  | Total | 80 | 1530667 |  |  | 1.000 |
|  | *Cladocopium C1* | Island | 10 | 240829 | 24082.9 | 13.0651 | 0.569 *** |
|  |  | Lineage | 3 | 52596 | 17532 | 9.5112 | 0.124 *** |
|  |  | Island:Lineage | 6 | 17676 | 2946 | 1.5982 | 0.042 ** |
|  |  | Residuals | 61 | 112442 | 1843.3 |  | 0.265 |
|  |  | Total | 80 | 423542 |  |  | 1.000 |

$p \leq 0.05$ \*; $p \leq 0.01$ \*\*; $p \leq 0.001$ \*\*\*

**Supplemental Table 11.** Permutational fitting of environmental variables to gene expression data. Table containing the results of permutational fitting of environmental variables to the gene expression data (vegan::envfit) in Pocillopora and Cladocopium C1 showing correlation coefficients (R^2^). Top contributing variables were those defined as having p ≤ 0.01 (*** and **).

| Variable | *Pocillopora* Host $R^2$ | *Cladocopium C1* Photosymbiont $R^2$ |
| --- | --- | --- |
| SST | 0.546 *** | 0.7131 *** |
| Primary Genetic Lineage (SVD Clade) | 0.5013 *** | 0.8765 *** |
| PO<sub>4</sub> | 0.4627 *** | 0.3795 *** |
| TSA DCW stdlength day | --- | 0.3477 *** |
| SST anomaly freq std (d) | 0.3626 *** | --- |
| TSA DHW lastrecovery (d) | 0.324 *** | 0.1087 |
| pH | 0.2823 *** | 0.3065 ** |
| TSA DCW minrecovery (d) | 0.2387 ** | 0.262 ** |
| TSA DCW stdlength (d) | 0.2339 ** | --- |
| chlorophyll | 0.2327 ** | 0.1078 |
| TSA heat max (°C) | 0.1808 ** | 0.2396 ** |
| PAR Sat | 0.1651 * | 0.3473 *** |
| TSA DHW lastduration (d) | 0.1458 * | 0.1895 * |
| TSA DCW lastrecovery (d) | 0.1441 * | 0.2545 ** |
| SiOH | 0.1343 * | 0.043 |
| SST anomaly max (°C) | 0.1237 * | 0.0476 |
| TSA DCW lastduration (d) | 0.0866 | 0.0443 |
| TSA cold freq std (d) | 0.077 | 0.023 |
| SST anomaly min (°C) | 0.0749 | --- |
| TSA DCW meanlength (d) | 0.0611 | 0.0717 |
| TSA DCW minlength (d) | 0.037 | 0.0222 |
| TSA cold min (°C) | --- | 0.0897 |
| TSA DCW meanrecovery day | --- | 0.0419 |

$p \leq 0.05$ \*; $p \leq 0.01$ \*\*; $p \leq 0.001$ \*\*\*

## Notes

### Competing Interest Statement

The authors have declared no competing interest.

